# *Caenorhabditis elegans* form sexually dimorphic and dynamic germ granules throughout meiotic prophase I progression

**DOI:** 10.64898/2026.04.27.721164

**Authors:** Acadia L. DiNardo, Diana E. Libuda

## Abstract

In sexually reproducing organisms, germ cells faithfully transmit both the genome and epigenetic information across generations through the formation of haploid gametes, such as eggs and sperm. Small RNA pathways tune gene expression in a sex-specific manner during germ cell development to facilitate both proper germ cell formation and transgenerational inheritance of epigenetic information. In *Caenorhabditis elegans*, components of small RNA pathways localize to germ granules, liquid-like membraneless organelles within the cytoplasm of developing germ cells. During oogenesis, germ granules form hierarchal sub-compartments that may be required for proper germ cell development and epigenetic inheritance. However, germ granule structure during spermatogenesis remains largely undescribed. Here we determine that the germ granule structural components PGL-1 and ZNFX-1 display sexually dimorphic foci morphology and size during meiotic prophase I progression. Further, we quantitate the sexually dimorphic sub-compartmentalization of these two proteins within the germ granule, determining that while PGL-1 and ZNFX-1 do associate during germ cell development, the extent of overlap varies between sexes and throughout meiotic progression. Additionally, we identify WAGO-4, a Argonaute protein central to gene regulation by small RNA pathways, as a sexually dimorphic component of the germ granule during germ cell development. Together, our studies reveal that the overall structure of the germ granule, as well as an Argonaute protein housed inside, are sexually dimorphic, which may underpin sex-specific regulation by small RNA pathways during germ cell development.

**Author Summary:** Small RNA pathways are critical for the regulation and passage of genomic and epigenetic information to the next generation. Components of these pathways are housed in germ granules during egg and sperm development. Previous work examining germ granule structure in *Caenorhabditis elegans* focused on oocytes and the late stages of meiosis I. Here, we comprehensively characterize the localization of two structural and one functional component of the germ granule throughout meiotic progression of both developing egg and sperm cells. We identify that both biophysical properties and germ granule configurations are dependent upon meiotic stage and sex.

## Introduction

Germ cell proliferation is critical for fertility and proper genome inheritance by the next generation. To faithfully transmit their genome from one generation to the next, sexually reproducing organisms create haploid germ cells, such as eggs and sperm, through the specialized cell division of meiosis. While the overall goals of chromosome pairing, recombination, and segregation are shared between egg and sperm development, the mechanisms that regulate gene expression and chromosome dynamics are highly dimorphic between oogenesis and spermatogenesis^19,47^. Indeed, a growing body of evidence suggests that oogenesis and spermatogenesis differ despite the conservation of numerous meiotic proteins^20,24,48,56,66,71^.

Gametes also transmit epigenetic information to the next generation. This form of inheritance, termed transgenerational epigenetic inheritance (TEI), is vital for preserving the immortality of the germline, as well as ensuring proper meiotic progression, fertilization, and embryogenesis^11,42^. Epigenetic information can be maintained between generations through mechanisms such as DNA methylation, histone modifications, and non-coding RNAs^36^. In humans and many other multicellular species, DNA methylation and histone marks undergo multiple rounds of reprograming and re-establishment during gametogenesis and early development^39,61,64,72^. Non-coding RNAs may serve as a molecular memory of some of these epigenetic DNA methylation and histone modification marks, as they provide a template for re-establishment of parental epigenetic states and directly target transcripts for silencing or licensing^7,9,18,54,76^. Small RNAs, such as PIWI-interacting RNAs (piRNAs) and endogenous small interfering RNAs (endo-siRNAs), are required for proper embryonic development and regulation of gene expression both transcriptionally and post-transcriptionally and are inherited through the cytoplasm of eggs^17,70,82^ and sperm^30,73,78^.

In *C. elegans,* small RNA pathways are crucial for TEI and regulate proper gene expression through generations utilizing an evolutionally well-conserved process known as RNA-interference (RNAi^35,51^). RNAi employs Argonaute proteins (AGOs) which bind small RNAs, 18 to 30 nucleotides in length, to target exogenous or endogenous sequences for post-transcriptional or co-transcriptional gene regulation^4,22,52,65,76,93^. Proper expression and regulation of these AGOs are required for meiotic progression, TEI, and stress response in germ cells^55,66,75,91,92^. Specifically, disruptions to AGOs and small RNA pathways cause sex-specific defects in gene regulation, passage and re-establishment of TEI, as well as fertility^10,74,75^. While these results suggest that AGOs and small RNA pathways can function in a sexually dimorphic manner, the mechanisms underlying these sexually dimorphic functions remain unknown.

During gametogenesis, several small RNA pathway components localize to germ granules, which are liquid-like phase-separated organelles found within the germ cells of most sexually reproducing organisms^32,38,68,83^. Germ granules are a type of biomolecular condensate whose formation is driven by biophysical interactions between RNA and multivalent proteins with intrinsically disordered domains^1,3,12,69^ and fine-tuned by concentrations of these macromolecules^8,32,79^. Condensates are hypothesized to act as sub-compartments within the cytoplasm or nucleus that function to mediate biochemical processes required for proper cellular functions^13,34,44,79^. Indeed, germ granules have been found to contain numerous components of small RNA pathways, and disruption to their localization and ability to phase separate during oogenesis and spermatogenesis has been tied to instances of infertility^21,81,89^. Foci morphology of phase separated condensates can act as a read out of the biophysical properties of the proteins and RNAs within itself. Moreover, larger sized foci are tied to higher protein concentrations and irregular aspherical surface area of foci can act as a proxy for more gel-like or glassy condensates^3,15,69,79^.

Previous work has found that while germ granules are primarily spherical, germ granules can change in shape due to localization around the nucleus or following fusion events^16,37^. In *C. elegans,* the germ granule is divided into sub-compartments with non-random configurations that are known to dynamically change in composition and localization through oogenesis and early germline development^25,45,58,60,85,87,91^. Currently, six distinct compartments of the germ granule have been identified, with two known to house the majority of AGOs associated with small RNA pathways and TEI^45,50,58,91^: 1) the P-granule, marked by the RNA-binding protein PGL-1^6,49,50^; and, 2) the Z-granule, which houses the zinc-finger protein ZNFX-1^46,91^. The P-granule contains multiple AGOs known to engage with endo-siRNAs, including WAGO-1 and WAGO-3, while the Z-granule includes a singular endo-siRNA interacting AGO, WAGO-4^40,45,73,91^.

While multiple AGOs, including WAGO-1, WAGO-3, and WAGO-4, are known to be incorporated into the germ granule during both oogenesis and spermatogenesis^40,73,74,77,91^, other AGOs are expressed, and therefore are a part of germ granule, only during oogenesis^77,90^ or spermatogenesis^23,28,29,63,77^. Additionally, previous work has found that one such constitutive AGO, WAGO-3, displays sexually dimorphic localization with germ granule components during late stages of meiosis^73^. Despite these characterized sexual dimorphisms in germ granule structure during meiotic progression, little is known how structural granule markers PGL-1 and ZNFX-1, as well as other constitutively expressed endo-siRNA interacting AGOs including WAGO-4 differs between the sexes. Indeed, most work characterizing the configuration of PGL-1 and ZNFX-1 within the germ granule, and how WAGO-4 co-localizes with them has been limited to presence versus absence of the proteins within sub-compartments during spermatogenesis^46,73,77,91^. Additionally, while previous studies have found that germ granule overlap and morphology is dynamic throughout gametogenesis, most work has focused on the localization of major sub-compartment components, such as PGL-1 and ZNFX-1, only during oogenesis^45,58,86,87,91^.

Here we examine germ granule dynamics through egg and sperm development by quantitatively assessing foci morphology and localization of the germ granule structural components PGL-1 and ZNFX-1. We find that the sub-compartments of the germ granule are sexually dimorphic in their morphology and localization relative to each other throughout germ cell development. Additionally, we identify WAGO-4, an Argonaute previously found to localize with ZNFX-1 during oogenesis, also displays sexually dimorphic foci localization and structure through germ cell development. Overall, our findings suggest that the germ granule organization is sexually dimorphic, which may contribute to sex-specific function of small RNA pathways.

## Methods and Materials

### Strains

All strains were generated for the N2 background and maintained at 20°C under standard conditions of nematode growth media (NGM) with OP50 *Escherichia coli.* All strains were kept as mating stocks, with all spermatogenesis specific work occurring in adult males 18-22 hours post-L4 stage and all oogenesis work occurring in adult hermaphrodites 18-22 hours post-L4 stage. The following strains were using in this study:

N2: Bristol wildtype strain
CB4108: CB4108: *fog-2(q71) V*.
DLW185: *wago-4(tm1019) II; mnC1 [dpy-10(e128) unc-52(e44) nls190 let?] II.* (This study)
YY916: *znfx-1(gg554[3xflag::gfp::znfx-1]) II*.
YY1325: *wago-4(gg620[3xflag::gfp::wago-4]) II*.

### WAGO-4 antibody production

Peptides within the IDR domain (RSEYQTSNDACIKRLEE-Cys) of the N-terminus of WAGO-4 were produced by Biomatik. Antibodies against the WAGO-4 peptides were produced in guinea pig and affinity purified by Pocono Rabbit Farms. Antibody specificity for WAGO-4 was confirmed utilizing immunofluorescence in a *wago-4(tm1019)* null mutant.

### Immunofluorescence

Immunofluorescence was performed as described in Claycomb *et al.,* 2009^26^ with the following adaptations. Briefly, gonads from adult male and hermaphrodite worms at 18-22 hour post-L4 stage were dissected together into 1x sperm salts (50 mM PIPES, pH 7.0, 25 mM KCl, 1 mM MgSO4, 45 mM NaCl, 2 mM CaCl2) on the same VWR Superfrost Plus slides, frozen and cracked on dry ice for 10 minutes and then fixed at -20°C for 5 minutes in 100% methanol, 5 minutes in 50% methanol/50% acetone, and 5 minutes in 100% acetone. Samples were blocked in 1xPBST + 3% BSA at room temperature for 15 minutes. Primary antibody dilutions were made in 1xPBST + 3% BSA and added to the slides. Slides were covered with a parafilm cover slip and incubated overnight at 4°C in a humid chamber. Slides were washed 3 x 10 minutes in 1xPBST and then blocked in 1xPBST + 3% BSA for 15 minutes at room temperature. Secondary antibody dilutions were made at 1:200 in 1x PBST + 3% BSA using Invitrogen goat Alexa Fluro-labeled antibodies and were added to the slides. Secondary antibody incubation occurred for 1 hour at room temperature in a dark humid chamber. Slides were then washed 3x10 minutes in 1xPBS in the dark and then incubated with 2 μg/ mL DAPI for 5 minutes in a humid chamber in the dark. Slides were then washed with 1xPBS for 5 minutes in the dark and then mounted with Vectashield. Slides were sealed with nail polish immediately following mounting and then stored at 4°C prior to imaging. All slides were imaged within 2 weeks of preparation. The following primary antibody dilutions were used: WAGO-4 anti-guinea pig (1:50, this study), rabbit anti-PGL-1 (1:1000, Susan Strome Lab), mouse anti-FLAG (1:1000, Millipore Sigma F1804-50UG).

### Imaging

Immunofluorescence slides were imaged at 1024 x 1024 pixel dimensions on an Applied Precision DeltaVision microscope with a 63x lens and a 1.5x optivar. Images were acquired as Z stacks at 0.2 μm intervals and deconvolved with Applied Precisions softWoRx deconvolution software.

### Foci quantification and overlap

Foci quantification adapted from Toraason *et al.,* 2021^84^. Briefly, germ granule foci were defined as Surface objects in Imaris (Bitplane) with the following settings: Smooth (not enabled), Background 0.513, and Seed Point Diameter (not enabled). Volume, sphericity, percent volume overlap, and X, Y, and Z data were exported from Imaris. Foci were analyzed if larger than 0.034 μm^2^ and smaller than 10 μm^2^ in volume.

Counts, volume, and sphericity of granule structures were then aligned along an X-Y axis utilizing a Gonad Linearization Algorithm as described in Toraason *et al.,* 2021^84^ and normalized from the pre-meiotic tip to the end of pachytene. Differences in counts were quantified using two sample Kolmogorov-Smirnov tests. Mean volume and sphericity were graphed using the plotnine.geom_smooth metric and the loess method along the length of the germline in a sliding window encompassing 0.1 units of normalized germline distance with a step size of 0.01 germline distance units. Mean values were calculated from foci analyzed in n=9 total germlines of each sex from ≥ 3 experimental replicates. Germlines were divided into 8 bins along meiotic progression utilizing well established nuclei morphology criteria to indicate meiotic stage: pre-meiotic tip (Bin 1), leptotene/zygotene (Bin 2), early pachytene (Bins 3-4), mid pachytene (Bins 5-6), late pachytene (Bins 7-8)^43^. Differences in distributions were calculated utilizing Mann-Whitney U test using SciPy stats.

To determine if foci are co-localized, data was binned as a binary, where 1 corresponded to foci that had a greater than 0% volume overlap with foci of interest and 0 corresponded to foci with 0% volume overlap with foci of interest. Foci from n=9 total germlines of each sex from ≥ 3 experimental replicates were analyzed and average proportion of foci with any overlap (value of 1) were graphed using the plotnine.geom_smooth metric and the loess method along the length of the germline in a sliding window encompassing 0.1 units of normalized germline distance with a step size of 0.01 germline distance units. Germlines were divided into 8 bins along meiotic progression utilizing well established nuclei morphology criteria to indicate meiotic stage: pre-meiotic tip (Bin 1), leptotene/zygotene (Bin 2), early pachytene (Bins 3-4), mid pachytene (Bins 5-6), late pachytene (Bins 7-8)^43^. Differences in probabilities between the sexes were calculated using Chi-squared test feature of SciPy stats.

Percent volume overlap was computed using foci that displayed any overlap with a granule component of interest (binary = 1). The percent volume overlap of each foci with corresponding protein of interest were then binned based upon meiotic stage utilizing well established nuclei morphology criteria to indicate meiotic stage: pre-meiotic tip (Bin 1), leptotene/zygotene (Bin 2), early pachytene (Bin 3), mid pachytene (Bin 4), late pachytene (Bin 5)^43^. Differences in distributions were calculated utilizing two sample Kolmogorov-Smirnov tests.

### Total RNA extraction and isolation for mRNA sequencing

300 adult hermaphrodites or adult males 18-22 hours post-L4 stage were picked onto unseeded plates and washed off with 1 mL of cold M9. Worms were then spun down for 1 minute at 2000 x g and placed on ice for 3 minutes. Supernatant was then aspirated off to not disturb the worm pellet and washed with 200 μL of M9 and again pelleted. Worms were washed one final time with 200 μL of DNase and RNase free UltraPure distilled water (Invitrogen), spun for 1 minute at 2000 RPM and all but 50uL of liquid was removed. 500 μL of TRIzol (ThermoFisher) was added and tubes were flash frozen and placed in -80°C freezer for 15 minutes. Samples were then removed and vortexed for 15 minutes at room temperature with 100 μm acid washed, RNase free beads (Sigma-Aldrich, <106 μm). Freeze-thaw and vortexing was then repeated twice more for a total of 3 times. 100 μL of chloroform was then added to samples and tubes were shaken vigorously for 15 seconds and incubated at room temperature for 3 minutes. Tubes were then centrifuged at 12,000 x g for 15 minutes at 4°C. The top aqueous layer was then removed and transferred to a new tube. Phenol: chloroform: isoamyl alcohol was added 1:1 in the new tube and mixed for 15 seconds. Samples were then centrifuged at 12,000 x g for 15 minutes at 4°C. Top aqueous phase was again transferred to a new tube and 20 μg of GlycoBlue (15 mg/mL, ThermoFisher) and 1:1 ratio of isopropanol was added to new tube. Samples were then incubated at -20°C overnight or -80°C for 1 hour. Samples were then centrifuged at 16,000 x g for 30 minutes at 4°C and supernatant then removed. The pellet was then with washed with 900 μL of 70% ice-cold ethanol for 10 minutes and then centrifuged at 16,000 x g for 10 minutes at 4°C. Supernatant was again removed and washing and centrifugation was repeated. Following last centrifugation, as much ethanol as possible was removed and pelleted was left to airdry for 10 minutes at room temperature. Pellet was then resuspended in 12 μL of RNAse-free water and purity and amount was quantified using Nanodrop. Samples were then diluted to 1 μg of RNA per 50 μL and used for downstream preparations.

### mRNA library preparation and sequencing

Samples were prepared for sequencing using the KAPA mRNA HyperPrep Kit for Illumina Platforms (KAPA Biosystems) following the protocol provided by manufacturer. The resulting DNA library was visualized and checked for quality using the 5200 Fragment Analyzer System (Agilent). Samples were then sequenced on the NovaSeq 6000 with a coverage of 25-30 million reads per a sample.

### mRNA-seq analysis

The mRNA sequences obtained from the sequencer were first assed for quality using MultiQC^33^. Adapter sequences were then removed using Trimmomatic for paired ends (version 0.36; Bolger et al., 2014) and run through MultiQC again to asses quality. The trimmed reads were then aligned to the *C.* elegans PRJNA13758 ce11 genome assembly (WormBase version WS280) using STAR (version 2.7.11b; Dobin et al., 2013). Reads were then counted using HTSeq (version 2.0.3; Anders et al., 2015) and differential expression was determined utilizing DESeq2 (version 3.20; Love et al., 2014). Genes were considered differentially expressed if their Log_2_ Fold Change was great than 1.58 or less than -1.58 to adjust for the presence of two gonads in hermaphrodites versus one in males and had a p-adjusted value of less than 0.05.

### Male and hermaphrodite fertility assays

For hermaphrodite fertility assays, hermaphrodites of each genotype were isolated at the L4 developmental stage and aged 18-22 hours to adult stage. Individual adult hermaphrodites were then singled on small plates (35 x 10 mm) with a small OP50 dot and allowed to lay for 24 hours. Following 24 hours, the adult hermaphrodite was moved to a fresh small plate (35 x 10 mm) with a small OP50 dot and allowed to lay for an additional 24 hours. This was repeated for a total of 5 plates. Following 24 hours on the fifth plate, the adult hermaphrodite was flamed off. 1-2 days following the removal of the adult hermaphrodite, plates were counted for number of living progeny, number of dead eggs, and number of unfertilized eggs. Plates where the hermaphrodite had run away or died were excluded. Differences in the ability of 3xFLAG::GFP::WAGO-4 hermaphrodites to produce viable progeny were analyzed using Two-way ANOVA with Tukey’s multiple comparison test.

For male fertility assays, males of each genotype and obligate females (CB4108: *fog2Δ*) obligate females were isolated at the L4 developmental stage and aged 18-22 hours to adult stage. Individual obligate females were then singled and paired with individual males on small plates (35 x 10 mm) with a small OP50 dot ringed with garlic extract to keep males from leaving the plate for 24 hours. After 24 hours males were moved to a new plate with an additional 18-22 hours post-L4 stage obligate female and allowed to mate for another 24 hours. After removal of the male, obligate females were allowed to lay eggs for 24 additional hours and then scored for living progeny, dead eggs, and unfertilized eggs. Pairs with no unfertilized eggs or where either the male or obligate female died before egg counting were excluded. Differences in the ability of 3xFLAG::GFP::WAGO-4 males to produce viable progeny were analyzed using Two-way ANOVA with Tukey’s multiple comparison test.

## Results

### Expression of germ granule components in *C. elegans* oocytes and spermatocytes

During germ cell development, germ granules house numerous proteins that regulate gene expression required for proper meiotic progression and fertility. A subset of these proteins are sexually dimorphic and known to only be expressed, and therefore incorporated, into the germ granule during either oogenesis or spermatogenesis^28,29,77,90^. Despite the sexually dimorphic composition of germ granules, numerous protein components are required for germ granule formation and localization during both oogenesis and spermatogenesis. PGL-1 and ZNFX-1 are two such proteins known to be fundamental components of germ granules independent of sex, forming distinct sub-compartments: the P-granule and Z-granule respectively^46,49,50,91^. Through mRNA-seq analysis of adult hermaphrodites undergoing oogenesis and adult males undergoing spermatogenesis (see Methods), we found that PGL-1 is upregulated in adult hermaphrodites compared to adult males, whereas ZNFX-1 expression does not significantly differ between the two sexes when corrected for gonad number (Fig. S1). To determine if these two constitutive germ granule components are sexually dimorphic in their localization and function, we characterized PGL-1 and ZNFX-1 foci morphology and co-localization during oogenesis and spermatogenesis.

### PGL-1 is a sexually dimorphic component of the germ granule

Using fixed immunofluorescence microscopy, we visualized and quantified PGL-1 localization in multiple whole germlines of wild-type adult hermaphrodites undergoing oogenesis and adult males undergoing spermatogenesis. As was previously observed^88^, PGL-1 is present around the nuclear periphery of developing germ cells throughout all of oogenesis (Fig. 1A, top) and through spermatogenesis until condensation into spermatids (Fig. 1A, bottom; Fig. S2A). Upon quantifying PGL-1 foci during meiotic prophase I progression in whole dissected germlines, we observed the recurrence of several sexually dimorphic trends of PGL-1 foci number across all wild-type germline datasets. From the pre-meiotic tip (PMT) through the mid-pachytene (MP) stages of meiosis I, we found that the average number of PGL-1 foci per a germ cell was largely the same (Fig. 1B). Upon reaching late pachytene (LP), however, developing oocytes nuclei had significantly more PGL-1 foci compared to stage-matched spermatocytes (Fig. 1B; oogenesis vs. spermatogenesis, Mann-Whitney U: LP 15.6 vs.12.3, p<0.032).

**Figure 1:**
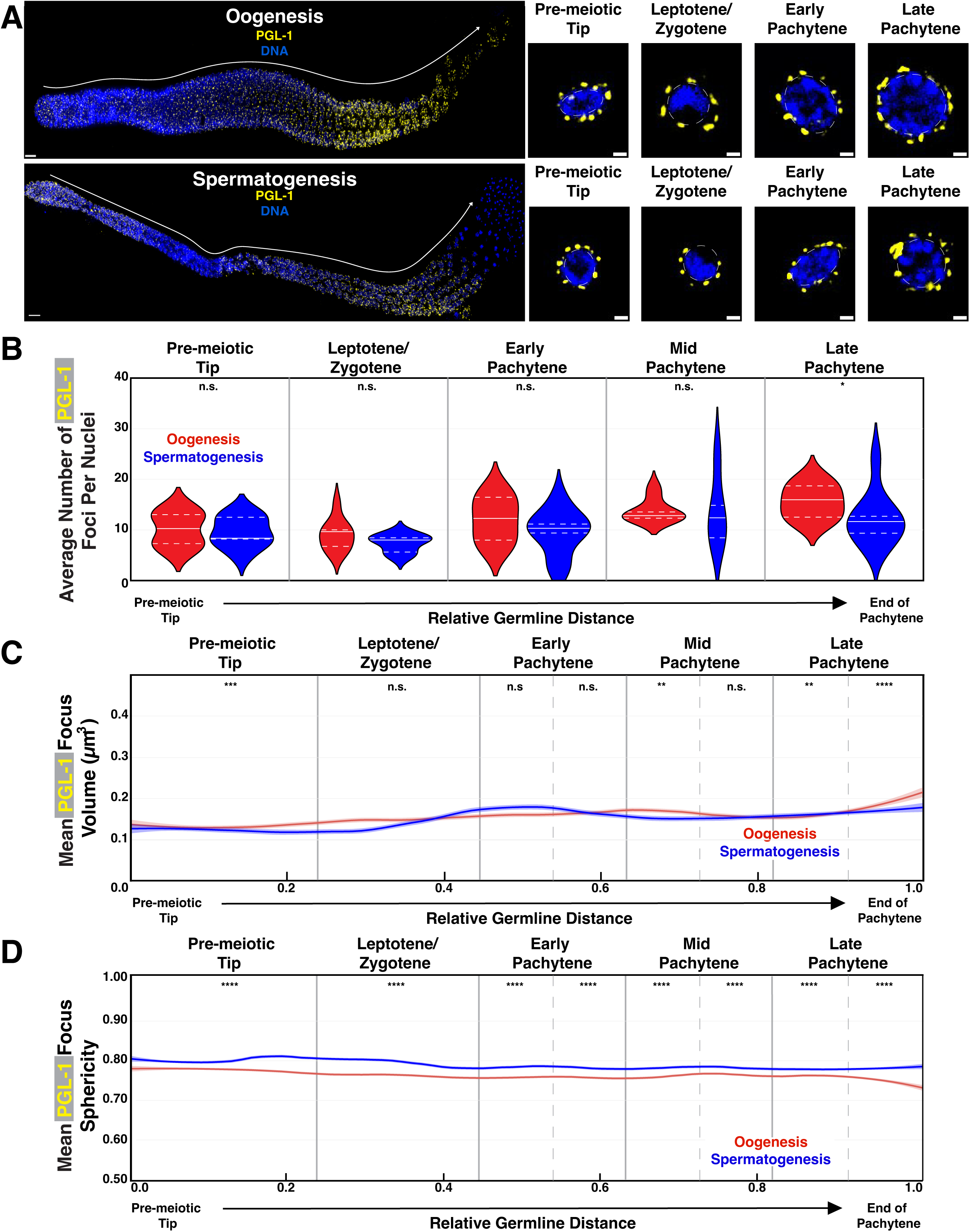
PGL-1 displays dynamic sexually dimorphic properties throughout meiosis prophase I progression. **(A)** Representative immunofluorescence images of germ granule structural protein PGL-1 (yellow) throughout meiosis of dissected germlines from wild-type adult hermaphrodite (oogenesis, top) and adult males (spermatogenesis, bottom). Enlarged panels display Z-slice of individual nuclei from four distinct stages of meiosis: pre-meiotic tip, leptotene/zygotene, early pachytene, and late pachytene. Scale bars represent 1 μm in insert panels and 10 μm in full gonad images. **(B)** Average number of PGL-1 foci per a nucleus during oogenesis (red) and spermatogenesis (blue). Nucleus and foci number per a region reported in Table S1 **(C)** Quantification of PGL-1 foci volume throughout oogenesis (red) and spermatogenesis (blue). Line plots represent the mean PGL-1 foci volume along the length of the germline calculated in a sliding window (*n*=9 gonads per sex; see Methods). Shaded areas around each line represents standard error of the mean. **(D)** Quantification of PGL-1 foci sphericity throughout oogenesis (red) and spermatogenesis (blue). Line plots represent the mean PGL-1 foci sphericity along the length of the germline are calculated in a sliding window (*n*=9 gonads per sex; see Methods). Shaded areas around each line represents standard error of the mean. For all statistical tests, p values were calculated using Mann-Whitney U tests. n.s. p>0.05, *p<0.05, **p<0.01, ***p<0.001, ****p<0.0001.

Our quantitative analysis of PGL-1 foci throughout meiotic progression revealed that PGL-1 focus volume is sexually dimorphic (Fig. 1C; Table S2; Table S3). While we observed that PGL-1 volume is similar from the onset of the PMT, we found that oocyte PGL-1 foci on average started to grow in volume in the latter portion of the PMT through the first part of leptotene/zygotene compared to spermatogenesis (Fig. 1C; Fig. S3A). In the second half of leptotene/zygotene, growth of PGL-1 foci in spermatogenesis led to similar volume size as observed in oogenesis which persisted through early pachytene (EP1 and EP2) (Fig. 1C; Fig. S3A). Finally, during the latter half of late pachytene (LP2), we found that PGL-1 foci volume increased in both sexes, albeit to a larger size during oogenesis vs spermatogenesis (Fig. 1C; Fig. S3A). Taken together, our data identifies quantitatively sexually dimorphic PGL-1 granule size that varies during meiotic prophase I progression.

To probe the biophysical properties of PGL-1 aggregates within the germ granule, we next interrogated the sphericity of PGL-1 foci between the sexes throughout meiotic progression. In previous works, sphericity of phase-separated compartments is correlated with liquid-like properties of the protein^2,3,16,37,59,62^. Compared to more spherical foci, which have more liquid-like protein properties with higher protein turnover, less spherical condensates tend to house proteins that are more gel-like in nature, leading to lower turnover with diffuse protein in the cytoplasm^3,16,27,37,53,62^. Throughout all stages of meiotic prophase I, we observed PGL-1 foci associated with nuclei undergoing spermatogenesis were significantly more spherical than those derived from oogenesis (Fig. 1D; Fig. S3B; oogenesis vs. spermatogenesis, Mann-Whitney U: PMT 0.770 vs. 0.794, p<0.0001; L/Z 0.756 vs. 0.787, p<0.0001; EP 0.751 vs. 0.776, p<0.0001; MP 0.756 vs. 0.775, p<0.0001; LP 0.750 vs. 0.773, p<0.0001). Together, our data indicates that PGL-1 is a sexually dimorphic component of the germ granule, displaying distinct foci size and morphological features dependent upon meiotic prophase I stage.

### ZNFX-1 granules display sexually dimorphic biophysical properties

We next analyzed the foci morphology and dynamics of ZNFX-1, a key component of the Z-granule sub-compartment, from adult hermaphrodites undergoing oogenesis and adult males undergoing spermatogenesis. As was found in several previous studies^46,87,91^, ZNFX-1 is perinuclear to developing oocytes and spermatocytes throughout the *C. elegans* germline (Fig. 2A). Our whole gonad imaging revealed that while PGL-1 foci rapidly grow diffuse in the condensation zone of spermatogenesis and are completely absent in mature spermatids (Fig. 1A, bottom; Fig. S2A), ZNFX-1 aggregates remain associated with developing spermatocytes late into the condensation zone (Fig. 2A, bottom), before forming large, non-nuclear associated foci present within the population of mature spermatids (Fig. S2B). Similar to PGL-1 foci, we found that in the PMT and L/Z regions of the germline the average number of ZNFX-1 foci per nucleus was not significantly different between oogenesis and spermatogenesis (Fig. 2B). Upon entering pachytene, we saw that the number of ZNFX-1 foci per nuclei increased specifically during oogenesis, but not spermatogenesis (Fig. 2B; oogenesis vs. spermatogenesis, Mann-Whitney U: EP 12.3 vs. 7.5, p<0.01). While we found no significant difference in MP between the sexes, by the end of pachytene there was again significantly more ZNFX-1 foci per a nucleus during oogenesis compared to spermatogenesis (Fig. 2B; oogenesis vs. spermatogenesis, Mann-Whitney U: MP 13.6 vs. 10.7, p=0.079; LP 10.1 vs. 8.2, p<0.001). These data suggest that the association of ZNFX-1 foci with the nuclei of developing germ cells is largely similar during early stages of germ cell development but sexually dimorphic during the early and late phases of pachytene in meiotic prophase I.

**Figure 2:**
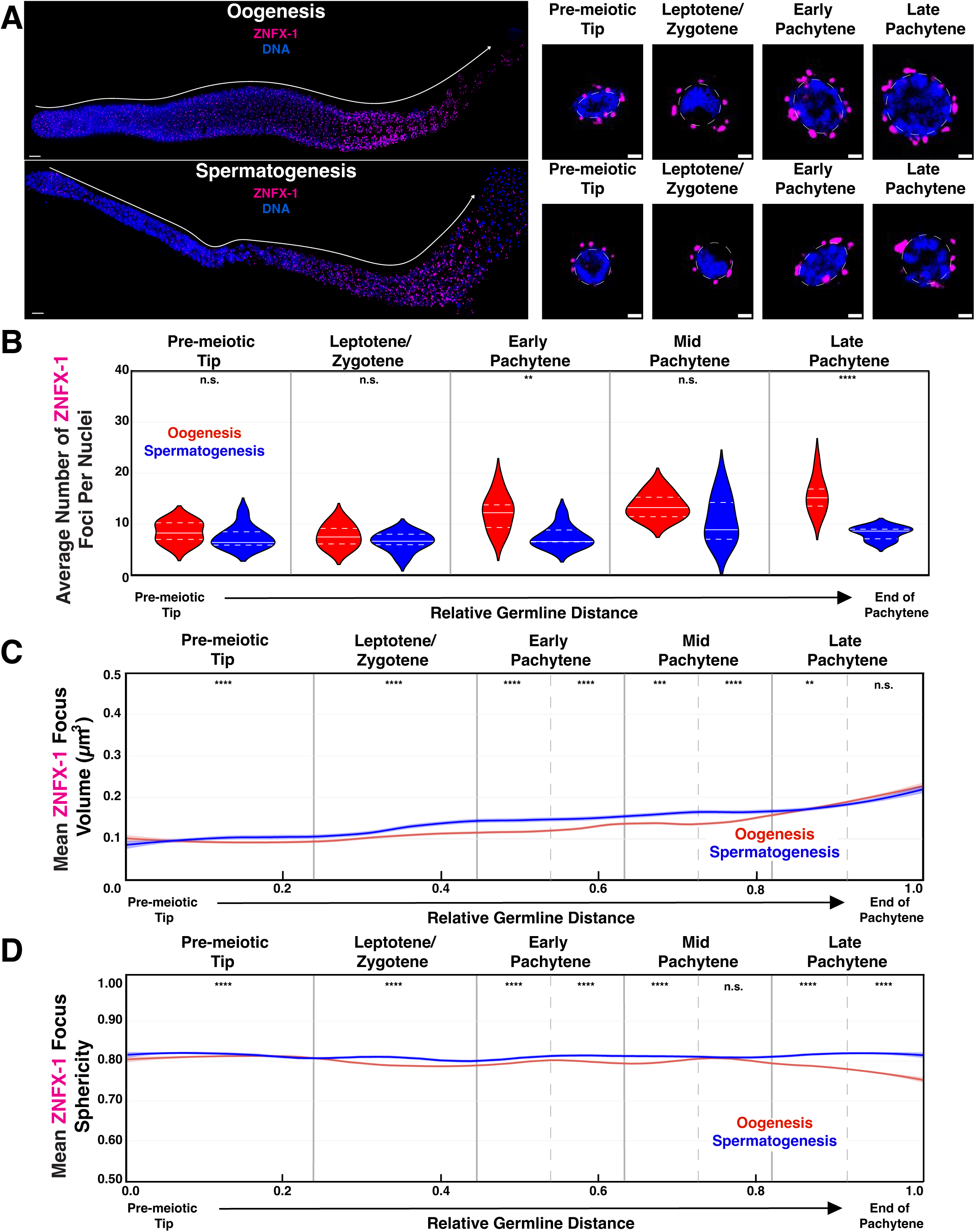
ZNFX-1 displays dynamic sexually dimorphic biophysical properties throughout meiosis. **(A)** Representative immunofluorescence images of germ granule structural protein ZNFX-1 (magenta) throughout meiosis of dissected germlines from wild-type adult hermaphrodite (oogenesis, top) and adult males (spermatogenesis, bottom). Enlarged panels display Z-slice of individual nuclei from four distinct stages of meiosis: pre-meiotic tip, leptotene/zygotene, early pachytene, and late pachytene. Scale bars represent 1 μm in insert panels and 10 μm in full gonad images. **(B)** Average number of ZNFX-1 foci per a nucleus during oogenesis (red) and spermatogenesis (blue). Nucleus and foci number per a region reported in Table S1 **(C)** Quantification of ZNFX-1 foci volume throughout oogenesis (red) and spermatogenesis (blue). Line plots represent the mean ZNFX-1 foci volume along the length of the germline calculated in a sliding window (*n*=9 gonads per sex; see Methods). Shaded areas around each line represents standard error of the mean. **(D)** Quantification of ZNFX-1 foci sphericity throughout oogenesis (red) and spermatogenesis (blue). Line plots represent the mean ZNFX-1 foci sphericity along the length of the germline are calculated in a sliding window (*n*=9 gonads per sex; see Methods). Shaded areas around each line represents standard error of the mean. For all statistical tests, p values were calculated using Mann-Whitney U tests. n.s. p>0.05, *p<0.05, **p<0.01, ***p<0.001, ****p<0.0001.

We next determined whether ZNFX-1 foci volume was also sexually dimorphic and dynamic through meiotic progression. While both PGL-1 and ZNFX-1 foci volume dynamically change through meiotic progression, the overall trends differed between the two components (Fig. 1C; Fig. 2C; Table S2; Table S3;). Unlike PGL-1, where foci volume largely did not differ between the sexes except in the distal and proximal ends of the germline (Fig. 1C), ZNFX-1 foci are significantly larger during spermatogenesis compared to oogenesis from PMT through MP (Fig. 2C; Fig. S3C; oogenesis vs. spermatogenesis, Mann-Whitney U: PMT 0.112μm^3^ vs. 0.120 μm^3^, p<0.0001; L/Z 0.130μm^3^ vs. 0.148μm^3^, p<0.0001; EP1 0.141μm^3^ vs. 0.175μm^3^, p<0.0001; EP2 0.155μm^3^ vs. 0182 μm^3^, p<0.0001; MP1 0.167μm^3^ vs. 0.192μm^3^, p<0.001; MP2 0.175μm^3^ vs. 0.206μm^3^, p<0.0001). Further, ZNFX-1 foci steadily grew in both sexes from PMT through MP (Fig. 2C; PMT vs MP2, Mann-Whitney U: oogenesis = 0.112μm^3^ vs. 0.175μm^3^, p<0.0001, spermatogenesis 0.120μm^3^ vs. 0.206μm^3^, p<0.0001). Prior to late pachytene, we found that ZNFX-1 foci constantly grew throughout meiotic progression for both sexes (Fig. 2C). Upon entrance to the late pachytene region of the germline (LP1), while ZNFX-1 foci associated with oocytes continued to grow in volume, ZNFX-1 foci largely plateaued in volume during spermatogenesis (Fig. 2C; MP2 vs. LP1, Mann-Whitney U: oogenesis = 0.175μm^3^ vs. 0.208μm^3^, p<0.0001; spermatogenesis = 0.206μm^3^ vs. 0.194μm^3^, p=0.182). By the end of late pachytene (LP2), though we found that ZNFX-1 foci again rapidly grew in both processes, leading to no significant difference in foci volume between oogenesis and spermatogenesis (Fig. 2C; oogenesis vs. spermatogenesis, Mann-Whitney U: LP2 0.252μm^3^ vs. 0.247μm^3^, p=0.120). These data suggest that, in addition to PGL-1, ZNFX-1 is a sexually dimorphic component of the germ granule that dynamically changes during meiotic prophase I progression. In contrast with PGL-1 foci, we observed a conserved trend between the sexes through the germline where ZNFX-1 foci grow in volume during both oogenesis and spermatogenesis.

To reveal sex-specific and meiotic-stage dependent differences in the biophysical environment, we also examined the sphericity of ZNFX-1 foci during oogenesis and spermatogenesis. Similar to PGL-1, we found that on average ZNFX-1 foci were largely more spherical during spermatogenesis compared to oogenesis from leptotene/zygotene through late pachytene (Fig. 2D; Fig. S3D; Table S2; Table S3). The notable exception is during the second half of mid pachytene where a decrease in sphericity during spermatogenesis coincides with an increase in ZNFX-1 foci sphericity during oogenesis leading to no significant difference in sphericity between the two processes (Fig. 2D; MP2 Mann Whitney U p=0.130). Together these data indicate that while meiotic stage may influence ZNFX-1 foci sphericity, the gamete sex may drive differences observed in sphericity.

### ZNFX-1 and PGL-1 co-localization is sexually dimorphic

To determine how ZNFX-1 and PGL-1 interface with each other throughout oogenesis and spermatogenesis, we analyzed the co-localization of these two proteins. In both sexes, we observed a significant proportion of PGL-1 and ZNFX-1 foci had some degree of overlap with each other throughout meiotic prophase I progression (Fig. 3A, white arrows). To better understand and quantify this overlap, we examined both the percentage of PGL-1 foci that overlapped with ZNFX-1 and the percentage of ZNFX-1 foci that had any overlap with PGL-1.

**Figure 3:**
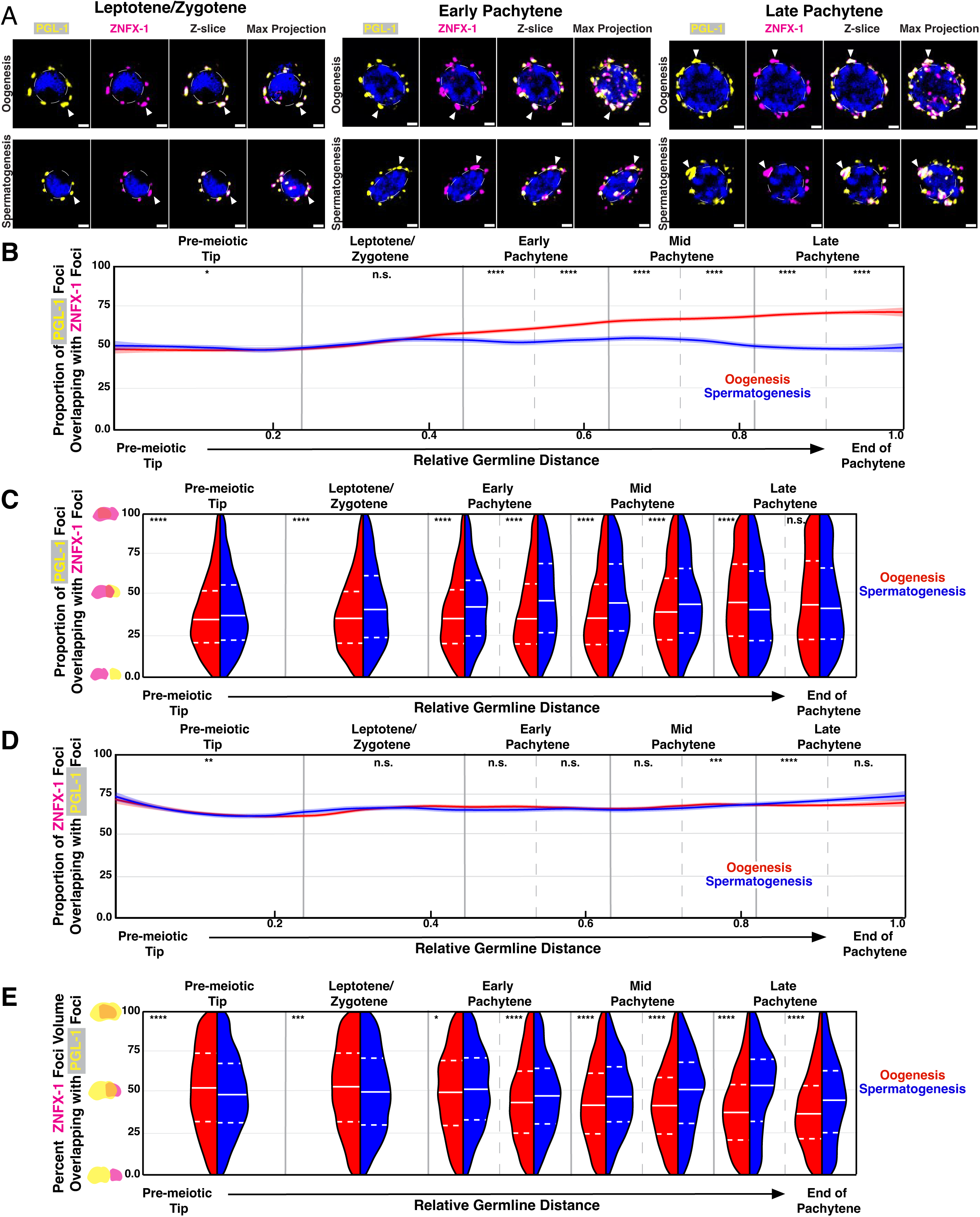
PGL-1 and ZNFX-1 co-localization is sexually dimorphic and dependent upon meiotic stage. **(A)** Representative images of single nuclei stained for PGL-1 (yellow) and ZNFX-1 (magenta) undergoing oogenesis (top) or spermatogenesis (bottom) during leptotene/zygotene (left), early pachytene (middle), and late pachytene (right). White arrows indicate examples of co-localization. Scale bars represent 1 μm. **(B)** Line plot representing the proportion of PGL-1 foci that overlap with ZNFX-1 along the length of the germline calculated in a sliding window (*n*=9 gonads per sex; see Methods). Shaded areas around each line represents standard error of the mean. **(C)** Split violin plot representing the distribution of PGL-1 volume overlap with ZNFX-1 in binned germline regions. **(D)** Line plot representing the proportion of ZNFX-1 foci that overlap with PGL-1 along the length of the germline calculated in a sliding window (*n*=9 gonads per sex; see Methods). Shaded areas around each line represents standard error of the mean. **E)** Split violin plot representing the distribution of ZNFX-1 volume overlap with PGL-1 in binned germline regions. For all split violin plots dashed white lines represent the first and third quartiles, solid white line represents the second quartile/median. For all statistical tests on binned germline regions of proportion overlapping, p values were calculated using Chi-squared tests. For all statistical tests on binned germline regions for percent volume overlap, p values were calculated using Mann Whiteny U test. n.s. p>0.05, *p<0.05, **p<0.01, ***p<0.001, ****p<0.0001.

We first looked at the proportion of PGL-1 foci that overlapped with ZNFX-1 during spermatogenesis compared to oogenesis throughout meiotic prophase I. Indeed, in agreement with our qualitative observations (Fig. 3A), around 50% of PGL-1 foci displayed some degree of overlap with ZNFX-1 throughout meiotic progression independent of sex (Fig. 3B). Similar to granule size and sphericity, we found that PGL-1 association with ZNFX-1 was dependent both upon process and meiotic stage. While PGL-1 foci overlap with ZNFX-1 foci was not significantly different between oogenesis and spermatogenesis from the PMT through L/Z, the proportion of PGL-1 associating foci progressively increased throughout all stages of pachytene during oogenesis (Fig. 3B; Table S4; Table S5). In contrast, PGL-1 foci associated with spermatocytes saw more varied changes in the proportion overlapping with ZNFX-1, leading to PGL-1 foci during spermatogenesis having significantly less overlap with ZNFX-1 compared to oogenesis during all stages of pachytene (Fig. 3B; oogenesis vs. spermatogenesis, Chi-squared: EP 63.0% vs. 54.4%, p<0.0001; MP 68.1% vs. 55.6%, p<0.0001; LP 71.9% vs. 49.8%, p<0.0001). Together, these data suggest that PGL-1 association with ZNFX-1 is dependent on sex, and that the magnitude of these differences in overlap is influenced by meiotic stage.

Previous works have identified that during oogenesis, ZNFX-1 forms distinct foci from PGL-1, suggesting that the Z- and P-granules function as separate sub-compartments from each other even when associated with the same germ granule^45,87,91^. To better capture instances where PGL-1 and ZNFX-1 may associate together within one granule but still form distinct sub-compartments, we analyzed the percent volume overlap between co-localized PGL-1 and ZNFX-1 foci. For all examined regions of the germline except the very end of pachytene (LP2), we found that the percent volume overlap of co-localized PGL-1 foci with ZNFX-1 was sexually dimorphic (Fig. 3C; Table S6; Table S7). From PMT through MP, PGL-1 foci from spermatogenesis had significantly more volume overlapping with ZNFX-1 foci compared to in oogenesis (Fig. 3C; mean volume overlap for oogenesis vs. spermatogenesis, Mann-Whitney U: PMT 38.4% vs. 41.0%, p<0.0001 L/Z 38.2% vs. 44.4%, p<0.0001; EP 39.3% vs. 46.5%, p<0.0001; MP 41.7% vs. 48.1%, p<0.0001). During the first half of late pachytene, we found a reversal in the trend observed through the rest of the germline where PGL-1 foci volume overlap with ZNFX-1 increased in oogenesis germ granules to be significantly higher than during spermatogenesis (Fig. 3C; LP1 47.5% vs. 43.9%, p<0.0001). In contrast, at the latter half of the late pachytene region, we found that the volume overlap of PGL-1 foci with ZNFX-1 were not significantly different between the oogenesis and spermatogenesis (Fig. 3C; LP2 47.1% vs. 46.1%, p=0.206). Taken together our data reveals that the composition and sub-compartmentalization of the germ granule is sexually dimorphic and highly dynamic during meiotic prophase I progression.

We next examined the reverse association: the proportion of ZNFX-1 foci overlapping with PGL-1 throughout oogenesis and spermatogenesis (Fig. 3D; Table S4; Table S5). We found that the association of ZNFX-1 with PGL-1 (magenta) was distinct from PGL-1 association with ZNFX-1 (yellow) independent of sex (Fig. S4; Table S8). From the PMT through EP during oogenesis (Fig. S4A) and throughout all of spermatogenesis (Fig. S4B), we found that significantly more ZNFX-1 foci overlapped with PGL-1 foci compared to PGL-1 foci associating with ZNFX-1. In both sexes, we found more than 63% of all ZNFX-1 foci associated with PGL-1, compared to some regions of the germline having less than 50% of PGL-1 foci localizing with ZNFX-1 (Fig. 3B; Fig 3D; Fig. S4). Despite these differences in proportion of PGL-1/ZNFX-1 association versus ZNFX-1/PGL-1 association observed within each sex, we found that ZNFX-1 foci localization with PGL-1 is largely similar between the sexes throughout meiotic prophase I (Fig. 3D). Together, these data indicate that the probability of association between PGL-1 and ZNFX-1 is dependent not only on sex, but also the protein of interest. Specifically, a higher proportion ZNFX-1 foci associate with PGL-1 compared to PGL-1 foci with ZNFX-1, suggesting that distinct populations of PGL-1 granules with and without ZNFX-1 may exist.

Finally, we determined whether the ZNFX-1 foci volume overlapping with PGL-1 was sexually dimorphic and dependent on meiotic stage (Fig. 3E; Table S6; Table S7). The median ZNFX-1 volume overlap with PGL-1 was indeed sexually dimorphic and dependent on meiotic stage. In early regions of the germline, including the premeiotic tip and leptotene/zygotene, we found that a larger percent of ZNFX-1 volume overlapped with PGL-1 during oogenesis (Fig. 3E; oogenesis vs. spermatogenesis, Mann-Whitney U: PMT 53.1% vs. 50.2%, p<0.0001, L/Z 53.3% vs. 51.5%, p<0.001). Upon entrance to the pachytene region of the germline, we found the inverse relationship: ZNFX-1 foci had more volume overlapping with PGL-1 during spermatogenesis compared to oogenesis (Fig. 3E; oogenesis vs. spermatogenesis, Mann-Whitney U: EP 47.9% vs. 51.0%, p<0.0001; MP 43.8% vs. 49.6%, p<0.0001; LP 39.6% vs. 48.6%, p<0.0001). These data indicate that even in regions of similar ZNFX-1 association with PGL-1 (Fig. 3D), there are sexual dimorphisms in the amount of overlap between these two granules (Fig. 3E). We find that PGL-1 more often forms a distinct granule from ZNFX-1, that is most prevalent during late meiosis I stages of spermatogenesis, while most ZNFX-1-containing granules associate with a PGL-1 focus. Despite the high frequency of PGL-1 and ZNFX-1 foci association, we found that there are still regions of the larger germ granule that only house PGL-1 or ZNFX-1. Overall, these data highlight the sexual dimorphic nature of ZNFX-1 and PGL-1 association with each other.

### WAGO-4 foci are sexually dimorphic through meiotic prophase I progression

The germ granule, and the sub-compartments marked by PGL-1 and ZNFX-1 are known to house numerous components of both endogenous and exogenous small RNA pathways. These components include the machinery required to make and amplify small RNAs, as well as Argonautes: proteins that bind small RNAs to aid in targeting of mRNAs for post-transcriptional gene regulation^45,52^. WAGO-4 is one such Argonaute that localizes specifically with ZNFX-1 and is known to act within multiple small RNA pathways, such as post-transcriptional gene regulation and transgenerational epigenetic inheritance (TEI) of silencing phenotypes^46,67,74,91,92^. Recent works have identified a sexually dimorphic role in WAGO-4-mediated TEI, where the AGO is expendable for TEI during spermatogenesis^74^. To better understand this sex-specific requirement for WAGO-4 in mediating transgenerational small RNA silencing, we examined if WAGO-4 behaves in a sexually dimorphic manner during meiotic prophase I. *wago-4* mRNA expression is sexually dimorphic, with an enrichment for *wago-4* transcripts in adult hermaphrodites undergoing oogenesis compared to adult males undergoing spermatogenesis when corrected for gonad number difference for each sex (Fig. S1). Additionally, we identified that tagging of the N-terminus of WAGO-4 causes male-specific fertility defects in brood size (Fig. S5), suggesting that while WAGO-4 may be present at higher levels in hermaphrodites versus males, proper WAGO-4 protein structure is imperative for preserving male fertility.

To determine if these sexually dimorphic phenotypes may also be associated with sexually dimorphic WAGO-4 localization, we utilized immunofluorescence microscopy to determine WAGO-4 localization during oogenesis and spermatogenesis (Fig. 4A). Due to our own and other reported observations of WAGO-4 tagging affecting fertility of adult hermaphrodites and males (Fig. S5^41^), we generated and confirmed the specificity of an antibody recognizing the N-terminus region of WAGO-4 (Fig. S6, see Methods). Using this antibody, we probed WAGO-4 accumulation, size, and morphology throughout multiple germlines undergoing oogenesis (Fig. 4A, top) and spermatogenesis (Fig. 4A, bottom). Similar to what we observed with PGL-1 and ZNFX-1, we found that the average number of WAGO-4 foci per nucleus was largely similar between oogenesis and spermatogenesis from the premeiotic tip through mid-pachytene (Fig. 4B). Upon late pachytene, there are significantly more WAGO-4 per nucleus during oogenesis compared to spermatogenesis (Fig. 4B; oogenesis vs. spermatogenesis, Mann-Whitney U: LP 15.7 vs. 11.0, p<0.01). The similar accumulation of more foci all three germ granule components suggests a shared need of more of these proteins localizing with developing eggs during late stages of meiosis I.

**Figure 4:**
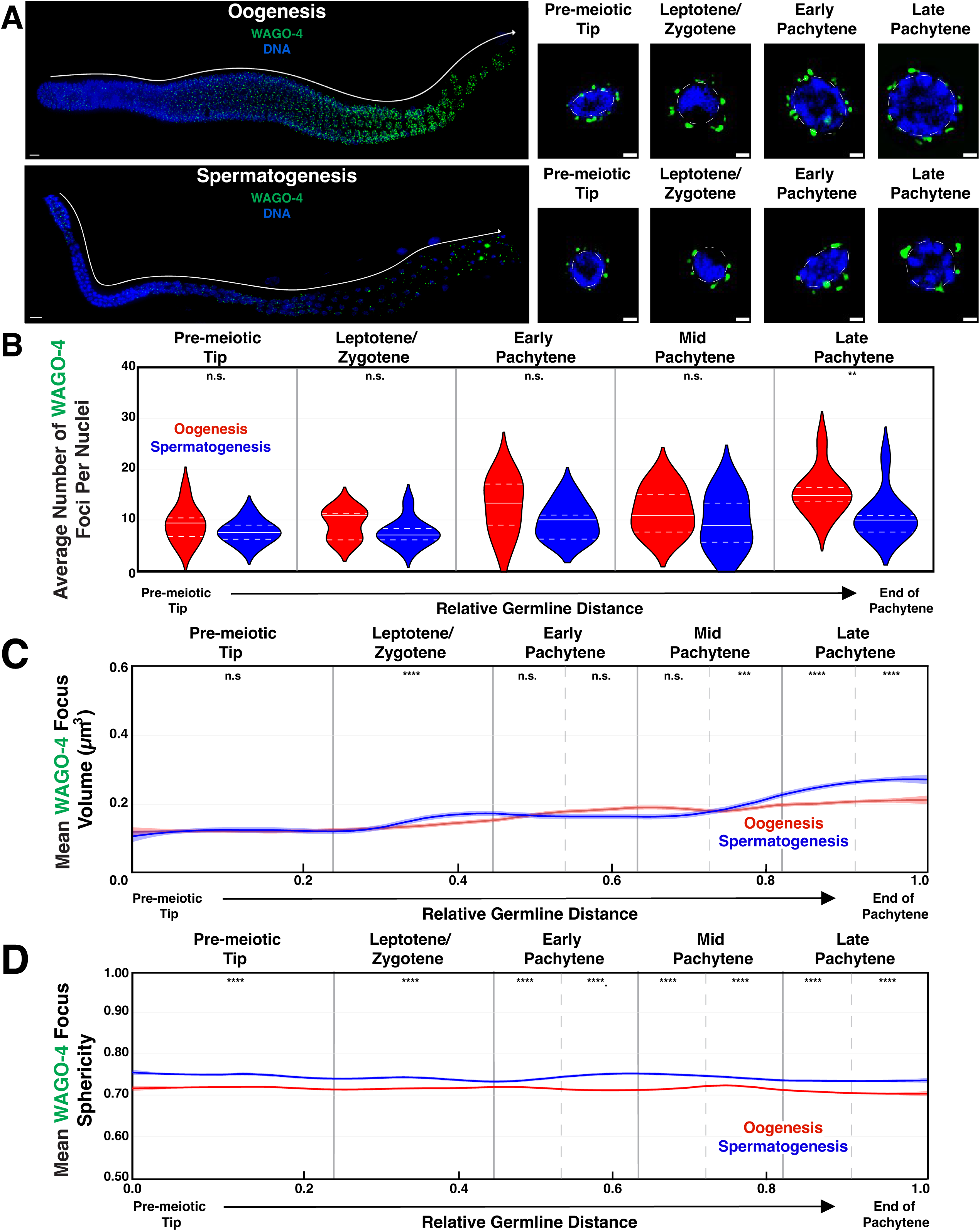
WAGO-4 displays dynamic sexually dimorphic biophysical properties throughout meiosis. **(A)** Representative immunofluorescence images of germ granule structural protein WAGO-4 (green) throughout meiosis of dissected germlines from wild-type adult hermaphrodite (oogenesis, top) and adult males (spermatogenesis, bottom). Enlarged panels display Z-slice of individual nuclei from four distinct stages of meiosis: pre-meiotic tip, leptotene/zygotene, early pachytene, and late pachytene. Scale bars represent 1 μm in insert panels and 10 μm in full gonad images. **(B)** Average number of WAGO-4 foci per a nucleus during oogenesis (red) and spermatogenesis (blue). Nucleus and foci number per a region reported in Table S1. **(C)** Quantification of WAGO-4 foci volume throughout oogenesis (red) and spermatogenesis (blue). Line plots represent the mean WAGO-4 foci volume along the length of the germline calculated in a sliding window (*n*=9 gonads per sex; see Methods). Shaded areas around each line represents standard error of the mean. **(D)** Quantification of WAGO-4 foci sphericity throughout oogenesis (red) and spermatogenesis (blue). Line plots represent the mean WAGO-4 foci sphericity along the length of the germline are calculated in a sliding window (*n*=9 gonads per sex; see Methods). Shaded areas around each line represents standard error of the mean. For all statistical tests on binned germline regions, p values were calculated using Mann-Whitney U tests. n.s. p>0.05, *p<0.05, **p<0.01, ***p<0.001, ****p<0.0001.

We next quantified the volume of WAGO-4 foci throughout meiotic prophase I progression for oogenesis and spermatogenesis (Fig. 4C; Fig. S7A; Table S2; Table S3). While there was no significant difference in the mean foci volume between the sexes in the PMT, WAGO-4 volume became significantly higher during spermatogenesis compared to oogenesis upon the onset of meiosis I with L/Z (Fig. 4C; Fig. S7A; oogenesis vs. spermatogenesis, Mann-Whitney U: L/Z 0.139μm^3^ vs. 0.148μm^3^, p<0.0001). Upon entrance into pachytene, WAGO-4 volume increased in oocytes, leading to WAGO-4 volumes no longer being sexually dimorphic for the first half of pachytene (Fig. 4C; Fig. S7A). The final half of pachytene exhibited an increase in WAGO-4 volume in both sexes, specifically with WAGO-4 foci growing significantly larger during spermatogenesis compared to oogenesis (Fig. 4C; Fig. S7A; oogenesis vs. spermatogenesis, Mann-Whitney U: MP2 0.193μm^3^ vs. 0.212μm^3^, p<0.001; LP1 0.199μm^3^ vs. 0.252μm^3^, p<0.0001; LP2 0.210μm^3^ vs. 0.268μm^3^, p<0.0001). To assess the biophysical properties of WAGO-4 aggerates during germ cell development, we quantified the changes in WAGO-4 sphericity throughout meiotic prophase I progression (Fig. 4D; Fig. S7B; Table S2; Table S3). In contrast with foci volume for which we identified dynamic changes in size between the sexes throughout meiotic progression, we found that WAGO-4 sphericity was significantly greater during spermatogenesis compared to oogenesis across the germline (Fig. 4D; Fig. S7B). Taken together, our data suggests that WAGO-4 is a dynamic component of the germ granule that displays distinct biophysical properties between sexes across germline. Further, we identified that changes in WAGO-4 volume between the sexes failed to correlate to changes in WAGO-4 sphericity, suggesting that foci morphology and size may be differentially regulated during germ cell development.

### WAGO-4 localization with PGL-1 changes through meiotic prophase I progression

Previous work found that WAGO-4 localizes and interacts with ZNFX-1 during oogenesis to coordinate transgenerational inheritance of gene regulation by small RNA pathways^91,92^. To determine if WAGO-4 displays sexually dimorphic co-localization with ZNFX-1 and PGL-1, we used immunofluorescence microscopy to stain for all three germ granule components in both sexes. We first determined the proportion of WAGO-4 foci that display any overlap with PGL-1 during oogenesis and spermatogenesis (Fig. 5A-B). Although we found no sexual dimorphisms in the proportion of WAGO-4 that associates with PGL-1 in the PMT (Fig. 5B; Table S4; Table S5; oogenesis vs. spermatogenesis, Chis-squared: PMT 69.6% vs. 68.4%, p=0.061), significantly more WAGO-4 foci associated with PGL-1 through the early stages of meiosis I during oogenesis compared to spermatogenesis (Fig. 5B; oogenesis vs. spermatogenesis, Chi-squared: L/Z 66.1% vs. 59.5%, p<0.0001; EP1 71.4 % vs. 51.2 %, p<0.0001; EP2 67.9% vs. 58.3%, p<0.0001; MP1 69.3% vs. 60.0%, p<0.0001). Through the latter half of pachytene, we found that WAGO-4 association with PGL-1 decreased during both oogenesis and spermatogenesis (Fig 5B; MP2 vs. LP2, Chi-squared: oogenesis 67.4% vs. 60.1%, p<0.0001; spermatogenesis 64.8% vs. 54.6%, p<0.0001). These data reveal that the proportion of WAGO-4 foci associated with PGL-1 varies between the gamete sexes and meiotic stages. Additionally, we identified two distinct populations of germ granules: granules where WAGO-4 associates with PGL-1 and granules where WAGO-4 and PGL-1 form distinct, independent foci from each other.

**Figure 5:**
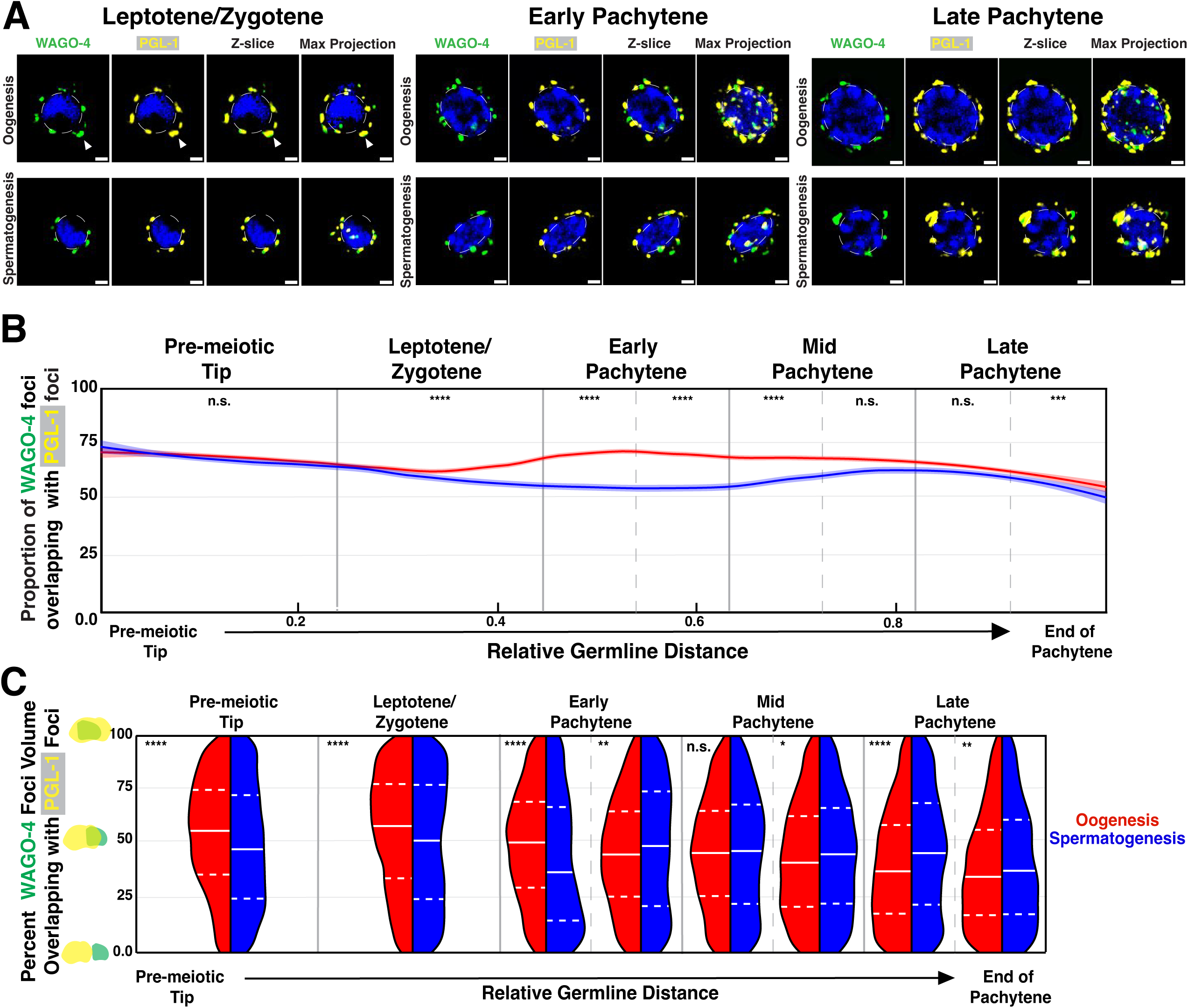
WAGO-4 co-localization with PGL-1 is sexually dimorphic and dynamic through meiotic progression. **(A)** Representative images of single nuclei stained for WAGO-4 (green) and PGL-1 (yellow) undergoing oogenesis (top) or spermatogenesis (bottom) during leptotene/zygotene (left), early pachytene (middle), and late pachytene (right). Scale bars represent 1 μm. **(B)** Line plot representing the proportion of WAGO-4 foci that overlap with PGL-1 along the length of the germline calculated in a sliding window (*n*=9 gonads per sex; see Methods). Shaded areas around each line represents standard error of the mean. **(C)** Split violin plot representing the differences in the distribution of WAGO-4 volume overlap with PGL-1 between oogenesis (red) and spermatogenesis (blue). Dashed white lines represent the first and third quartiles, solid white line represents the second quartile/median. For all statistical tests on binned germline regions of proportion overlapping, p values were calculated using Chi-squared tests. For all statistical tests on binned germline regions for percent volume overlap, p values were calculated using Mann Whiteny U test. n.s. p>0.05, *p<0.05, **p<0.01, ***p<0.001, ****p<0.0001.

To determine if the population of PGL-1-associated WAGO-4 foci display significant co-localization with PGL-1, and how these proportions of overlap between PGL-1 and WAGO-4 foci vary between egg and sperm development, we examined the percent volume overlap between WAGO-4 foci that co-localize with PGL-1 (Fig. 5C; Table S6; Table S7). We found that when only examining WAGO-4 foci that associate with PGL-1, the percent volume overlap was dependent both on sex and meiotic stage. From the PMT until the first half of early pachytene (EP1), we saw significantly higher WAGO-4 percent overlap volume with PGL-1 during oogenesis compared to spermatogenesis (Fig. 5C; oogenesis vs. spermatogenesis, Mann-Whitney U: PMT 54.6% vs. 49.1%, p<0.0001; L/Z 55.7% vs. 50.9%, p<0.0001; EP1 50% vs. 42.5%, p<0.0001). From the second of half of early pachytene (EP2) and largely through the end of late pachytene we observed the reciprocal was true, where more WAGO-4 foci volume overlapped with PGL-1 during spermatogenesis compared to oogenesis (Fig. 5C; oogenesis vs. spermatogenesis, Mann-Whitney U; EP2 45.7% vs. 49.0%, p<0.01; MP2 43.2% vs. 45.5%, p<0.05; LP1 39.8% vs. 46.5%, p<0.0001; LP2 38.6% vs. 41.4%, p<0.01). Taken together, these data indicate that while most WAGO-4 foci display association with PGL-1, the majority of WAGO-4 granule volume fails to co-localize with PGL-1 foci.

### WAGO-4 foci are sexually dimorphic in co-localization dynamics with ZNFX-1

We next examined how WAGO-4 interfaces with the Z-granule marker, ZNFX-1, throughout meiotic prophase I progression. WAGO-4 is a key component of the Z-granule during oogenesis, with mutations in WAGO-4 mirroring a number of phenotypes observed in ZNFX-1 mutant contexts during oogenesis^67,91^. Throughout meiotic prophase I, more than half of the WAGO-4 foci co-localize with ZNFX-1 independent of sex and meiotic stage (Fig. 6 A-B). Similar to what we observed with WAGO-4 co-localization with PGL-1, there are sexually dimorphic regions of the germline where significantly more WAGO-4 foci localize with ZNFX-1 during oogenesis compared to spermatogenesis (Fig. 6B; Table S4; Table S5). Interestingly, while we observed this sexually dimorphic association of WAGO-4 with both PGL-1 and ZNFX-1 within the leptotene/zygotene through the first half of mid pachytene, WAGO-4 only displayed sexually dimorphic association with ZNFX-1 within the PMT (Fig. 6B; oogenesis vs. spermatogenesis, Chi-squared: PMT 63.3% vs. 57.2%, p<0.0001). Together, these data indicate that WAGO-4 association with ZNFX-1 and their proportional overlap is sexually dimorphic and dependent on meiotic prophase I stage.

**Figure 6:**
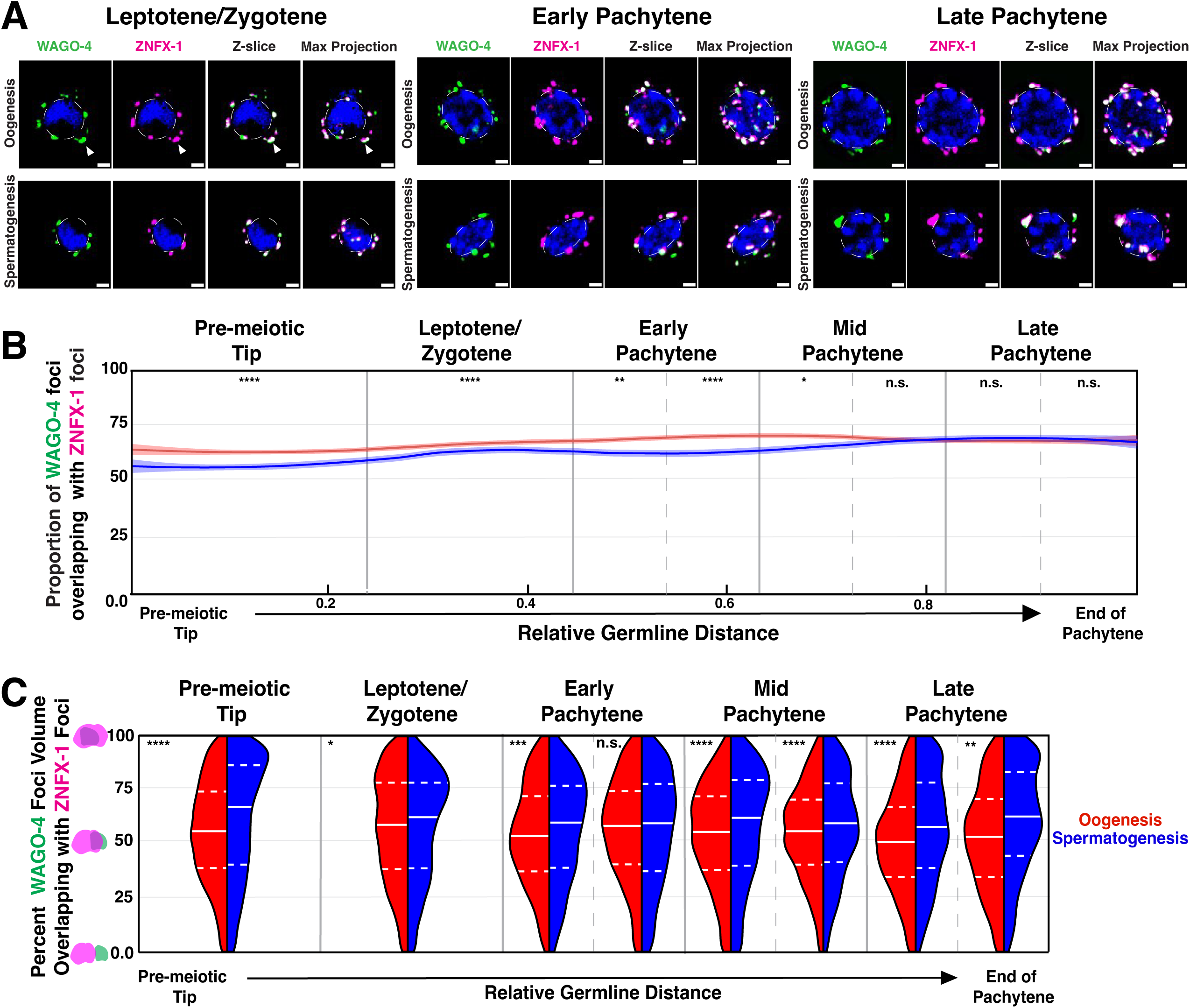
WAGO-4 co-localization with ZNFX-1 is sexually dimorphic and dynamic through meiotic progression. **(A)** Representative images of single nuclei stained for WAGO-4 (green) and ZNFX-1 (magenta) undergoing oogenesis (top) or spermatogenesis (bottom) during leptotene/zygotene (left), early pachytene (middle), and late pachytene (right). Scale bars represent 1 μm. **(B)** Line plot representing the proportion of WAGO-4 foci that overlap with ZNFX-1 along the length of the germline calculated in a sliding window (*n*=9 gonads per sex; see Methods). Shaded areas around each line represents standard error of the mean. **(C)** Split violin plot representing the differences in the distribution of WAGO-4 volume overlap with ZNFX-1 between oogenesis (red) and spermatogenesis (blue). Dashed white lines represent the first and third quartiles, solid white line represents the second quartile/median. For all statistical tests on binned germline regions of proportion overlapping, p values were calculated using Chi-squared tests. For all statistical tests on binned germline regions for percent volume overlap, p values were calculated using Mann Whiteny U test. n.s. p>0.05, *p<0.05, **p<0.01, ***p<0.001, ****p<0.0001.

To determine whether there are changes in the extent of co-localization of WAGO-4 with ZNFX-1 between sexes, we examined the percent volume overlap for each WAGO-4 focus with ZNFX-1 (Fig. 6C; Table S6; Table S7). Through almost all stages of meiotic progression, WAGO-4 foci had significantly higher volume overlap with ZNFX-1 during spermatogenesis compared to oogenesis (Fig. 6C; oogenesis vs. spermatogenesis, Mann-Whitney U: PMT 55.7% vs. 62.1%, p<0.0001; EP1 54.1% vs. 57.0%, p<0.001; MP1 54.1% vs. 58.1%, p<0.0001; MP2 55.0% vs. 58.5%, p<0.0001; LP1 50.8% vs. 58.0%, p<0.0001; LP2 52.8% vs. 61.7%, p<0.01). Taken together, these data indicate that WAGO-4 foci more often localize to the Z-granule during oogenesis, suggesting a sexually dimorphic role and perhaps function for WAGO-4. Furthermore, despite higher WAGO-4 association with ZNFX-1 during oogenesis, WAGO-4 foci that do associate with ZNFX-1 during spermatogenesis demonstrate a larger percent of co-localization compared to oogenesis, suggesting a difference in total interface between the two proteins within individual germ granules.

### WAGO-4 P-granule and Z-granule association is sexually dimorphic

Based on our observations that a large proportion of WAGO-4 foci overlap with both PGL-1 and ZNFX-1, and that the overlapping foci volume is substantial, we next wanted to compare the relative WAGO-4 interfaces with both structural proteins. To this end, we concurrently examined WAGO-4 localization with both germ granule structural proteins throughout meiotic prophase I progression (Fig. 7A). During oogenesis, we found that the proportion of WAGO-4 overlapping with ZNFX-1 (Fig. 7B, magenta; Table S6; Table S8) and PGL-1 (Fig. 7B, yellow; Table S6; Table S8) is dependent upon meiotic stage. In the PMT during oogenesis, we found that a larger proportion of WAGO-4 foci associates with PGL-1 compared to ZNFX-1 (Fig. 7B; WAGO-4/PGL-1 vs. WAGO-4/ZNFX-1, Chi-squared: PMT 69.9% vs. 63.3%, p<0.0001). An increase in WAGO-4/ZNFX-1 localization and a concurrent decrease in WAGO-4/PGL-1 association upon entrance into L/Z leads to no significant difference in WAGO-4 localization with both granule components (Fig. 7B; WAGO-4/PGL-1 vs. WAGO-4/ZNFX-1, Chi-squared: L/Z 66.1% vs. 66.9%, p=0.182). Through pachytene, local increases and decreases in WAGO-4 localization with each structural component drive highly variable WAGO-4 association with both proteins. During the first half of early pachytene(EP1), we found higher WAGO-4 localization with PGL-1, while the second half of early pachytene(EP2) and all of late pachytene saw higher WAGO-4 association with ZNFX-1, and finally mid-pachytene had no significant difference in WAGO-4 association with either structural component (Fig. 7B; WAGO-4/PGL-1 vs. WAGO-4/ZNFX-1, Chi-squared: EP1 71.4% vs. 68.9%, p<0.05; EP2 67.9% vs. 72.8%, p<0.0001; MP 68.3% vs. 68.7%, p=0.639; LP 62.1% vs. 67.8%, p<0.0001). In contrast during spermatogenesis, we found that WAGO-4 predominantly associated with each structural protein in distinct regions of the germline. In the PMT of spermatogenesis, we found that a higher proportion of WAGO-4 foci associated with PGL-1 (Fig. 7C; WAGO-4/PGL-1 vs. WAGO-4/ZNFX-1: PMT 68.4% vs. 57.2). Upon entry L/Z, and persisting through the reminder of spermatogenesis, a higher proportion of WAGO-4 foci localized with ZNFX-1 compared to PGL-1 (Fig. 7C: WAGO-4/PGL-1 vs. WAGO-4/ZNFX-1, Chi-squared; L/Z 59.5% vs. 62.1%, p<0.01; EP 54.7% vs. 63.9%, p<0.0001; MP 62.4% vs. 67.9%, p<0.0001; LP 58.6% vs. 68.9%, p<0.0001). This change in association is driven by an increase throughout meiotic prophase I progression of the WAGO-4/ZNFX-1 association, while WAGO-4 localization with PGL-1 follows an inverse trend (Fig. 7C; L/Z vs. LP2, Chi-squared; WAGO-4/PGL-1 59.5% vs. 54.6%, p<0.01; WAGO-4/ZNFX-1 62.1% vs. 68.3%, p<0.0001). Together, these data indicate that during spermatogenesis, the proportion of WAGO-4 association with each germ granule component is driven by meiotic stage, where ZNFX-1 association increases while PGL-1 decreases. During oogenesis, we see a more varied pattern of WAGO-4 foci association, suggesting that the co-localization of WAGO-4 with just one of these key germ granule components may be less important for proper germ cell development in oogenesis versus spermatogenesis.

**Figure 7:**
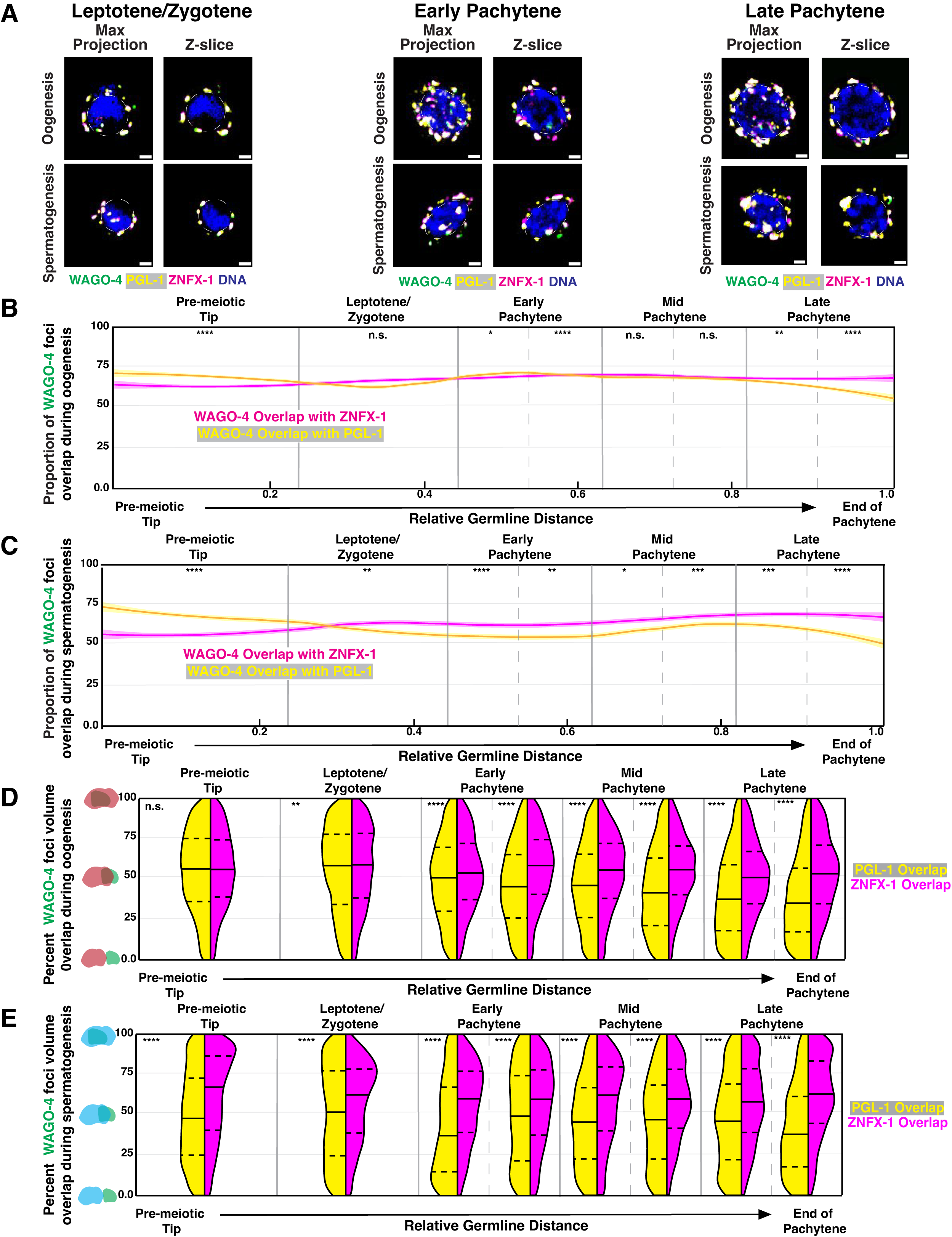
WAGO-4 displays protein specific co-localization patterns during germ cell development. **(A)** Representative images of WAGO-4 (green), PGL-1 (yellow), and ZNFX-1 (magenta) localization around single nuclei undergoing oogenesis (top) and spermatogenesis (bottom) during (left) leptotene/zygotene, (middle) early pachytene, and (right) late pachytene. **(B)** Line plot representing the proportion of WAGO-4 foci that overlap with PGL-1 (yellow) and ZNFX-1 (magenta) during oogenesis along the length of the germline calculated in a sliding window (*n*=9 gonads per sex; see Methods). Shaded areas around each line represents standard error of the mean. **(C)** Line plot representing the proportion of WAGO-4 foci that overlap with PGL-1 (yellow) and ZNFX-1 (magenta) during spermatogenesis along the length of the germline calculated in a sliding window (*n*=9 gonads per sex; see Methods). Shaded areas around each line represents standard error of the mean. **(D)** Split violin plot representing the differences in the distribution of WAGO-4 volume overlap between ZNFX-1 (magenta) and PGL-1 (yellow) during oogenesis. **(E)** Split violin plot representing the differences in the distribution of WAGO-4 volume overlap between ZNFX-1 (magenta) and PGL-1 (yellow) during spermatogenesis. On split violin plots, dashed black lines represent the first and third quartiles, solid black line represents the second quartile/median. For all statistical tests on binned germline regions of proportion overlapping, p values were calculated using Chi-squared tests. For all statistical tests on binned germline regions for percent volume overlap, p values were calculated using Mann Whiteny U test. n.s. p>0.05, *p<0.05, **p<0.01, ***p<0.001, ****p<0.0001.

Finally, to better understand how WAGO-4 foci that localize with PGL-1 and ZNFX-1 may be sub-compartmentalized with both proteins when present in the same granule, we compared the percent volume overlap of WAGO-4 with ZNFX-1 and PGL-1 during both oogenesis (Fig. 7D; Table S9) and spermatogenesis (Fig. 7E; Table S9). During oogenesis, we found that while in the distal region of the germline, WAGO-4 overlaps relatively equally with both PGL-1 and ZNFX-1 (Fig. 7D; WAGO-4/PGL-1 vs. WAGO-4/ZNFX-1, Mann-Whitney U: PMT 54.6% vs. 55.7%, p=0.066), upon entrance into meiosis I, WAGO-4 displayed significantly more overlap with ZNFX-1 compared to PGL-1 (Fig. 7D; WAGO-4/PGL-1 vs. WAGO-4/ZNFX-1, Mann-Whitney U: L/Z 55.7% vs. 57.3%, p<0.01; EP 48% vs. 55.4%, p<0.0001; MP 44.7% vs. 54.6%, p<0.0001; LP 39.3% vs. 51.8%, p<0.0001). In contrast, throughout all of stages of spermatogenesis, WAGO-4 foci had significantly more volume overlap with ZNFX-1 compared to PGL-1 (Fig. 7E; WAGO-4/PGL-1 vs. WAGO-4/ZNFX-1, Mann-Whitney U: PMT 49.1% vs. 62.1%, p<0.0001; L/Z 50.9% vs. 58%, p<0.0001; EP 45.9% vs. 56.8%, p<0.0001; MP 45.9% vs. 58.3%, p<0.0001; LP 43.8% vs. 60.1%, p<0.0001). Together, these data indicate that WAGO-4 more frequently localizes with ZNFX-1, and therefore the Z-granule, over PGL-1 throughout all stages of spermatogenesis and following entrance into meiosis I during oogenesis.

## Discussion

### Sexual dimorphisms within the small RNA pathway

Our study reveals that ZNFX-1 and PGL-1 have sexually dimorphic foci morphology and co-localization throughout germ cell development. Previous studies established that PGL-1 and ZNFX-1 play distinct roles from each other during germ cell development. ZNFX-1 is required for proper transgenerational epigenetic silencing of transcripts through its association with small RNA pathway machinery^46,91^. Indeed, ZNFX-1 has been found to maintain TEI by storing pUGylated transcripts, which are utilized as templates for secondary sRNA transcription^80^ within the germ granule, as well as preventing drift of secondary sRNA transcription from the 3’ end to the 5’ end of templates^46,67^. In contrast, proper PGL-1 structure is required for AGO-specific mediated regulation of transcripts within the germline and silencing of spermatogenic genes during oogenesis^6,21,81,89^. The majority of previous research describing germline specific functions of ZNFX-1 and PGL-1 predominately studied adult hermaphrodites undergoing oogenesis^6,21,46,91^; however, numerous meiotic proteins function differently during oogenesis versus spermatogenesis to mediate conserved processes including chromosome pairing and recombination^20,56,71^. Our identification of differences in ZNFX-1 and PGL-1 localization within the germ granule support hypotheses that mechanisms of TEI and gene silencing may be mediated by core components of the germ granule, including WAGO-4^74^. To better determine whether functional roles of ZNFX-1 and PGL-1 are conserved between the sexes, there is a growing need for comprehensive sex-comparative studies of both ZNFX-1 and PGL-1.

In addition to TEI and AGO-mediated transcript regulation, both ZNFX-1 and PGL-1 directly interact with multiple AGOs required for functional gene regulation during germ cell development^6,40,46,73,91^. Sexually dimorphic AGO gene expression during meiosis is well documented in *C. elegans.* Numerous AGOs (including ALG-3/4, ERGO-1, VSRA-1, and WAGO-10) are expressed only during oogenesis or spermatogenesis^23,28,29,63,77,90^. Moreover, constitutively expressed AGOs (including PRG-1 and WAGO-3) have sex-specific roles or localization patterns during pachytene^66,73^. Here, we demonstrate that although WAGO-4 is endogenously expressed during both oogenesis and spermatogenesis, it exhibits sexually dimorphic features including foci size and sphericity during meiosis I. Notably, we find that WAGO-4 foci are on more spherical during spermatogenesis compared to oogenesis, which indicates WAGO-4 likely has more liquid-like properties during spermatogenesis and more gel-like properties during oogenesis (Fig. 4D). Additionally, we found that sub-compartmentalization of WAGO-4 was distinct between the sexes despite significantly more WAGO-4 foci volume overlapping more with ZNFX-1 compared to PGL-1 during both spermatogenesis and oogenesis (Fig. 7). These data suggests that WAGO-4 may play sex-specific roles in regulating TEI and piRNA-directed silencing through interactions with PGL-1 and ZNFX-1^46,73,77,91^. Further work probing the functional impacts of sex-specific WAGO-4 localization and biophysical properties will improve our understanding of how the mechanisms of TEI and piRNA-mediated gene silencing promote proper germ cell development and fertility.

### Meiosis I stage-specific function of small RNA pathways

Our work highlights not only sexually dimorphic foci morphology and localization of PGL-1, ZNFX-1, and WAGO-4, but also that these foci are dynamic throughout meiotic prophase I progression. During both spermatogenesis and oogenesis, we find that the characteristics and interactions of each focus differs throughout meiotic progression. Indeed, ZNFX-1 foci progressively grow in volume across the germline independent of sex (Fig. 2C), while PGL-1 and WAGO-4 experience local regions of growth and loss of volume dependent on meiotic stage. These differences in foci behavior, despite all examined proteins being constitutive components of the germ granule during meiosis, may underscore the varying protein-protein or protein-RNA interactions that occur throughout meiotic progression within the germ granule. This hypothesis is further supported by our observations in oogenesis that there is a higher incident of ZNFX-1-independent PGL-1 foci in early meiotic prophase I in comparison to mid- and late stages of pachytene (Fig. S4A). The higher instance of ZNFX-1-independent PGL-1 foci only occurring within the PMT through EP could suggest a specific function for PGL-1 that is independent of ZNFX-1 that is not required during later stages of meiosis I during oogenesis.

We also find that WAGO-4 displays dynamic changes throughout meiotic progression between its localization with both PGL-1 and ZNFX-1. With PGL-1, we find that WAGO-4 association and percent overlap with the protein generally decreases throughout meiotic progression (Fig. 5B; Fig. 5C). By contrast, WAGO-4 association with ZNFX-1 increases (Fig. 6B; Fig, 6C). These responses suggests that WAGO-4 function, as well as the interacting proteins and RNAs, may change dynamically over the course of meiotic prophase I, potentially in a sexually dimorphic fashion. Future work parsing specific WAGO-4 interactions during distinct stages of meiotic progression, and how their disruption changes the transcriptional and translational landscape, will greatly aid in our understanding of stage specific roles of small RNA mediated regulation.

Together our work uncovers novel sexually dimorphic morphology and localization pattens of both constitutive germ granule components and a functional AGO during germ cell development. These differences in germ granule sub-compartmentalization highlight a new avenue of study in understanding mechanisms of small RNA pathway regulation and transmission of epigenetic information during oogenesis and spermatogenesis.

## Data Availability

The mRNA-sequencing data set generated in this study are available on the NCBI BioProject database (https://www.ncbi.nlm.nih.gov/bioproject/) under accession number PRJNA1252786. Strains and WAGO-4 antibodies are available upon request.

## Acknowledgments

We thank Z. Bush, J. Brown, C. Crahan, C. Cahoon, N. Kurhanewicz, and H. Wilson for thoughtful discussion and comments on the manuscript. We also thank A. Barkan, B. Bowerman, D. Garcia, and S. Hansen for feedback on figures. We thank the Kennedy lab for sharing the 3xFLAG::GFP::ZNFX-1 and the 3xFLAG::GFP:WAGO-4 strains and the Strome Lab for sharing their PGL-1 α rabbit antibody. We are grateful to the University of Oregon’s Genomics and Cell Characterization Core Facility for mRNA sample prep and sequencing. We thank the CGC for providing multiple strains for this study (funded by National Institutes of Health P40 OD010440).

## Funding Sources

This work was supported by the National Institutes of Health T32GM007431 to A.L.D. and National Institutes of Health R35GM128890 to D.E.L.

## Conflict of Interest

The authors declare no conflicts of interest.

## Supplementary Material

**Supplemental Figure 1:**
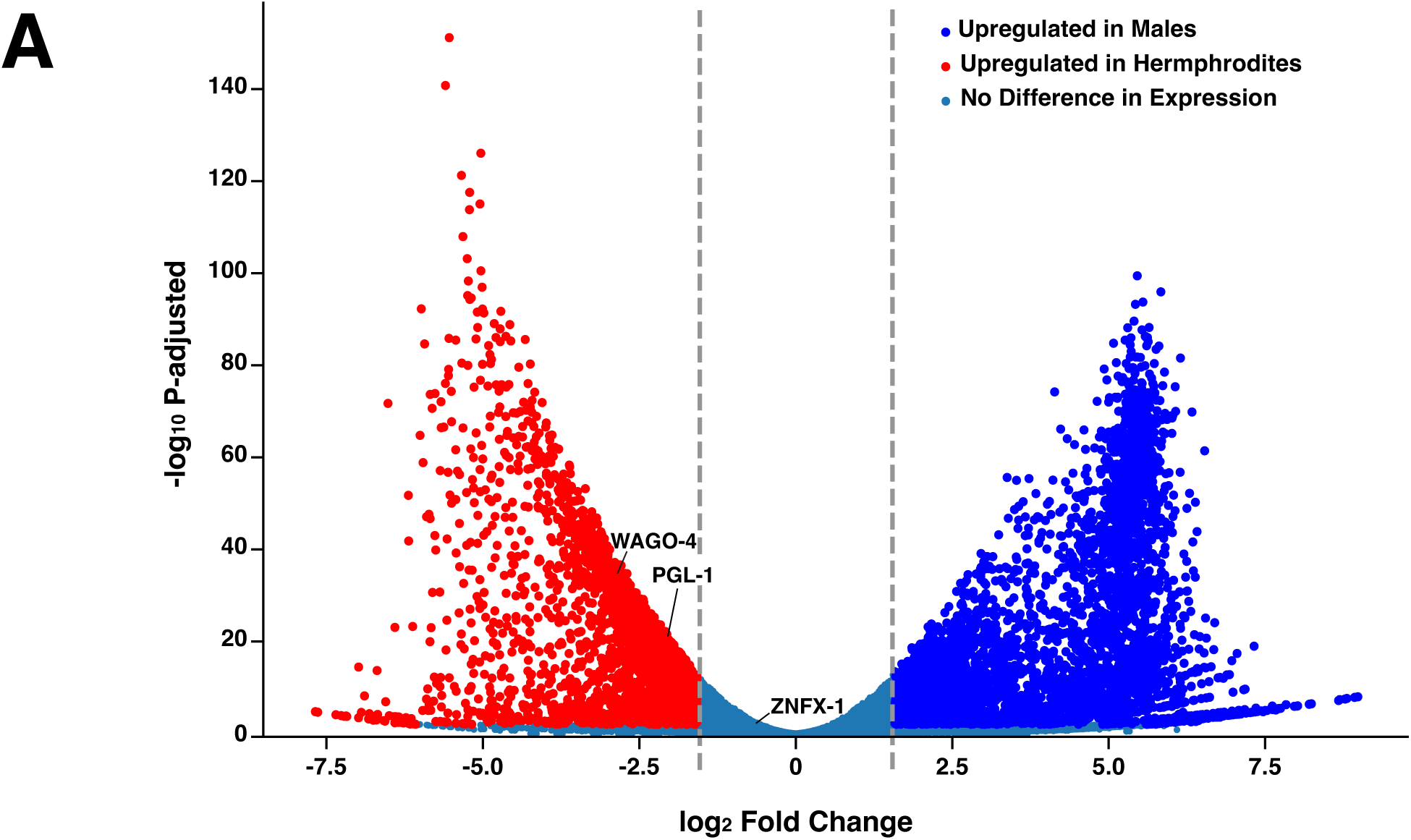
WAGO-4 and PGL-1 display sexually dimorphic gene expression. **(A)** Volcano plot of differentially expressed genes from mRNA-seq data between wild-type adult males (spermatogenesis) and adult hermaphrodites (oogenesis). Genes upregulated in hermaphrodites compared to males are in red. Genes upregulated in males compared to hermaphrodites are in blue. Blue-grey dots represent genes with no statistical difference in regulation between the sexes. Genes with significant fold changes had a log_2_ fold change larger than ±1.58 and a p-value < 0.05. Labels denote genes of interest further explored in this study.

**Supplemental Figure 2:**
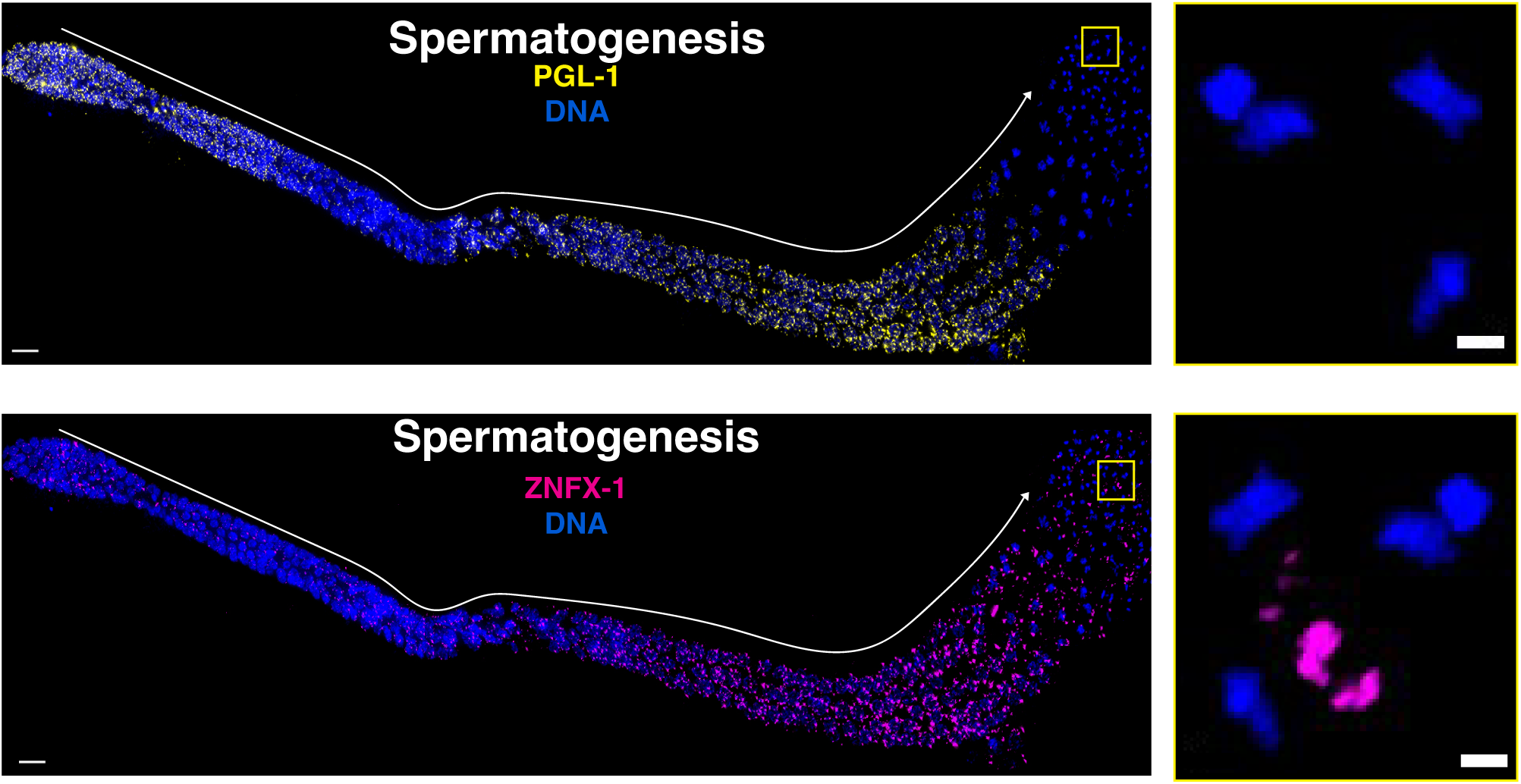
PGL-1 and ZNFX-1 foci localization with condensed spermatids. **(A)** Representative immunofluorescence images of germ granule structural protein PGL-1 (yellow) throughout meiosis of dissected germlines from wild-type adult males (spermatogenesis). Enlarged panel display max projection of condensed sperm nuclei. Scale bars represent 1 μm in insert panels and 10 μm in full gonad images. **(B)** Representative immunofluorescence images of germ granule structural protein ZNFX-1 (magenta) throughout meiosis of dissected germlines from wild-type adult males (spermatogenesis). Enlarged panel display max projection of condensed sperm nuclei. Scale bars represent 1 μm in insert panels and 10 μm in full gonad images.

**Supplemental Figure 3:**
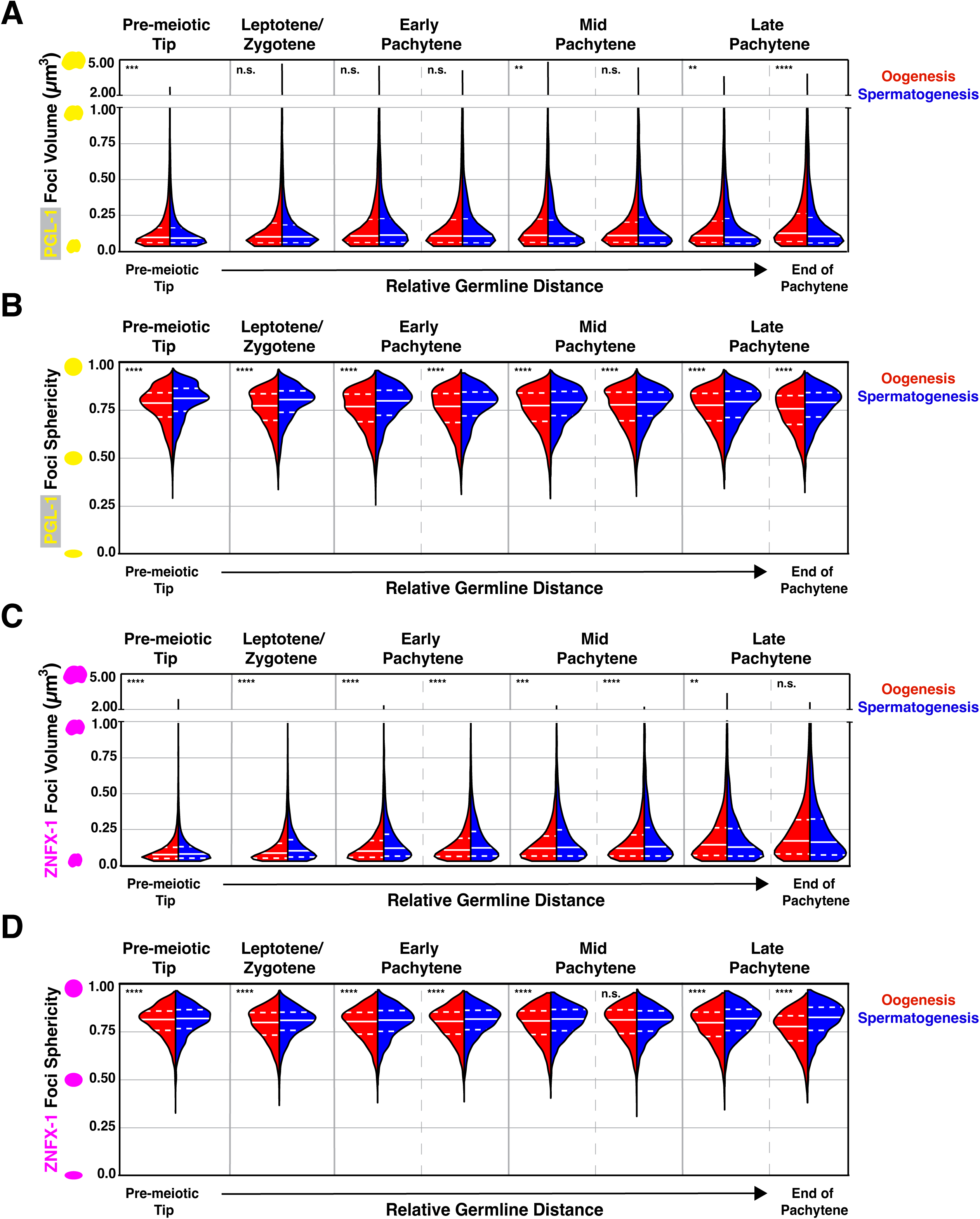
Volume and sphericity distribution of PGL-1 and ZNFX-1 foci during germ cell development. **(A)** Split violin plot depicting the volume distribution of PGL-1 foci during oogenesis (red) and spermatogenesis (blue). **(B)** Split violin plot depicting the sphericity distribution of PGL-1 foci during oogenesis (red) and spermatogenesis (blue). **C)** Split violin plot depicting the volume distribution of ZNFX-1 foci during oogenesis (red) and spermatogenesis (blue). **(D)** Split violin plot depicting the sphericity distribution of ZNFX-1 foci during oogenesis (red) and spermatogenesis (blue). Dashed white lines represent the first and third quartiles, solid white line represents the second quartile/median. For all statistical tests, p-values were calculated using two-sample Kolmogorov-Smirnov test. n.s. p>0.05, *p<0.05, **p<0.01, ***p<0.001, ****p<0.0001.

**Supplemental Figure 4:**
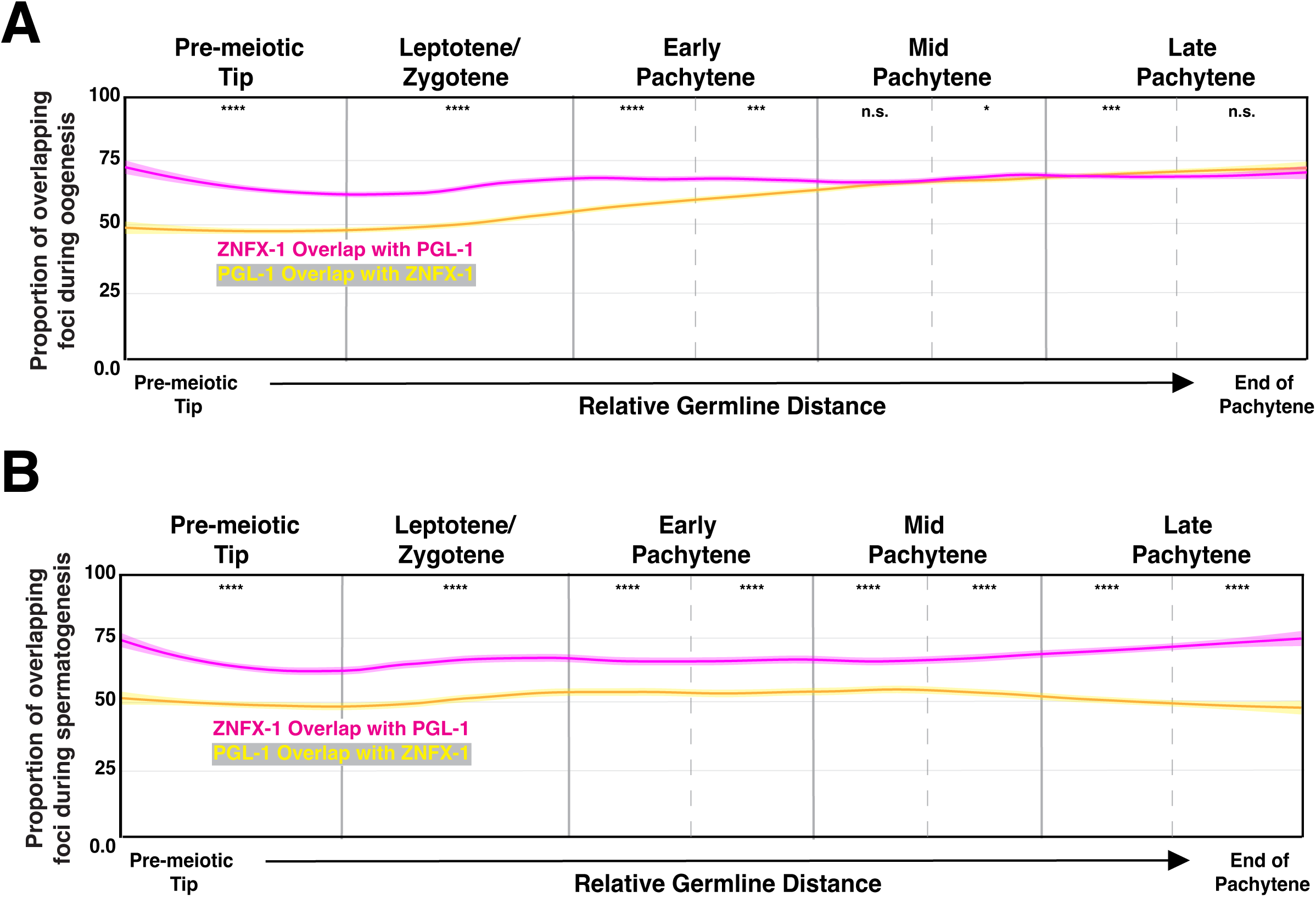
Overlap of ZNFX-1 and PGL-1 during oogenesis and spermatogenesis. **(A)** Line plot representing the proportion of ZNFX-1 foci that overlap with PGL-1 (magenta) and PGL-1 overlap with ZNFX-1 (yellow) during oogenesis along the length of the germline calculated in a sliding window (*n*=9 gonads per sex; see Methods). Shaded areas around each line represents standard error of the mean. **(B)** Line plot representing the proportion of ZNFX-1 foci that overlap with PGL-1 (magenta) and PGL-1 overlap with ZNFX-1 (yellow) during spermatogenesis along the length of the germline calculated in a sliding window (*n*=9 gonads per sex; see Methods). Shaded areas around each line represents standard error of the mean. For all statistical tests on binned germline regions, p values were calculated using Chi-squared tests. n.s. p>0.05, *p<0.05, **p<0.01, ***p<0.001, ****p<0.0001.

**Supplemental Figure 5:**
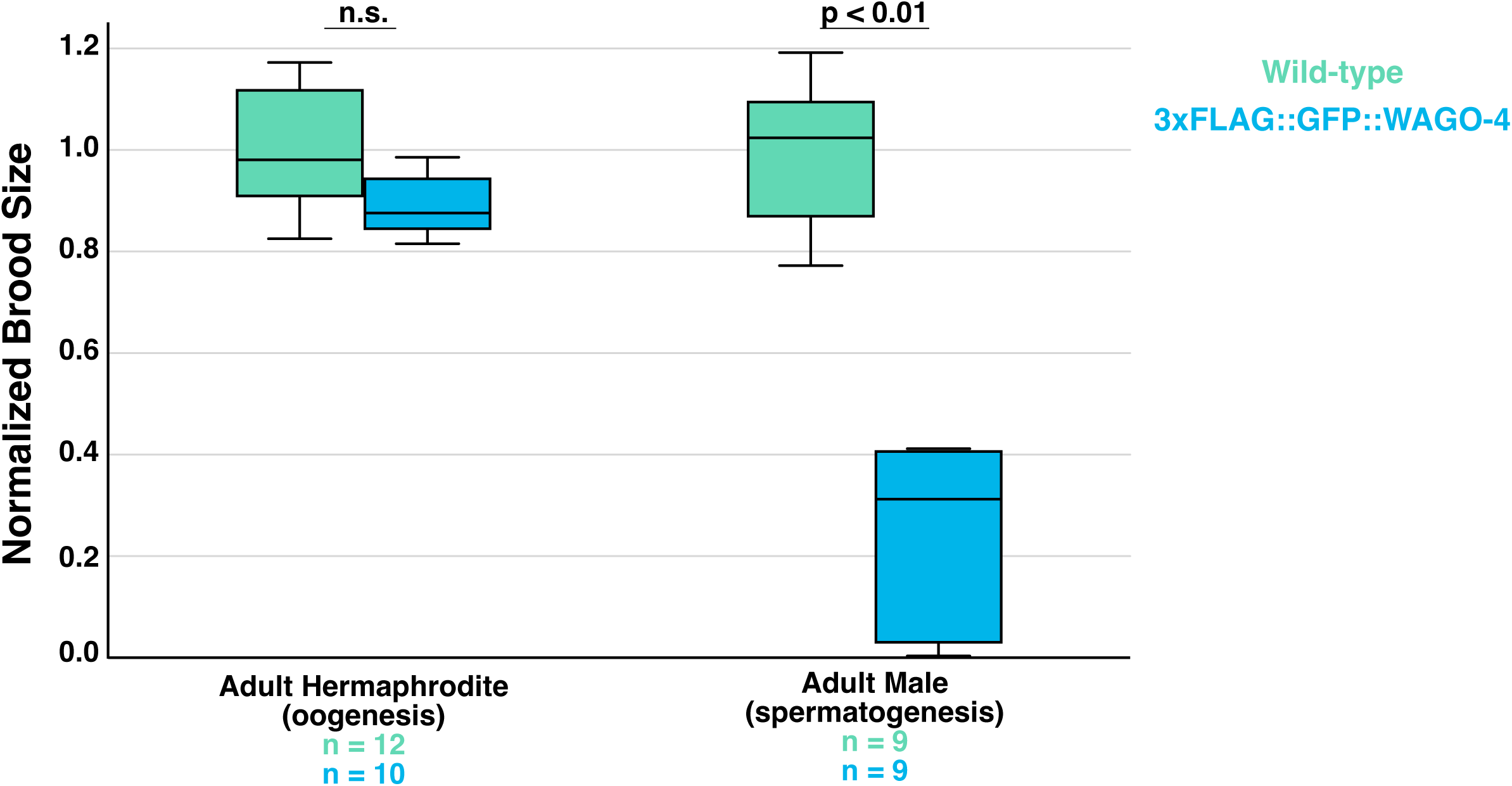
Tagging of WAGO-4 N-terminus causes significant decrease in male brood size. Normalized brood size of adult hermaphrodites undergoing oogenesis (left) and adult males undergoing spermatogenesis (right) from wild-type (light green) and endogenously tagged 3xFLAG::GFP::WAGO-4 *C. elegans.* Brood sizes include any dead embryos but exclude unfertilized eggs. Reported n values indicate number of *C. elegans* analyzed. P-values calculated using One-way ANOVA.

**Supplemental Figure 6:**
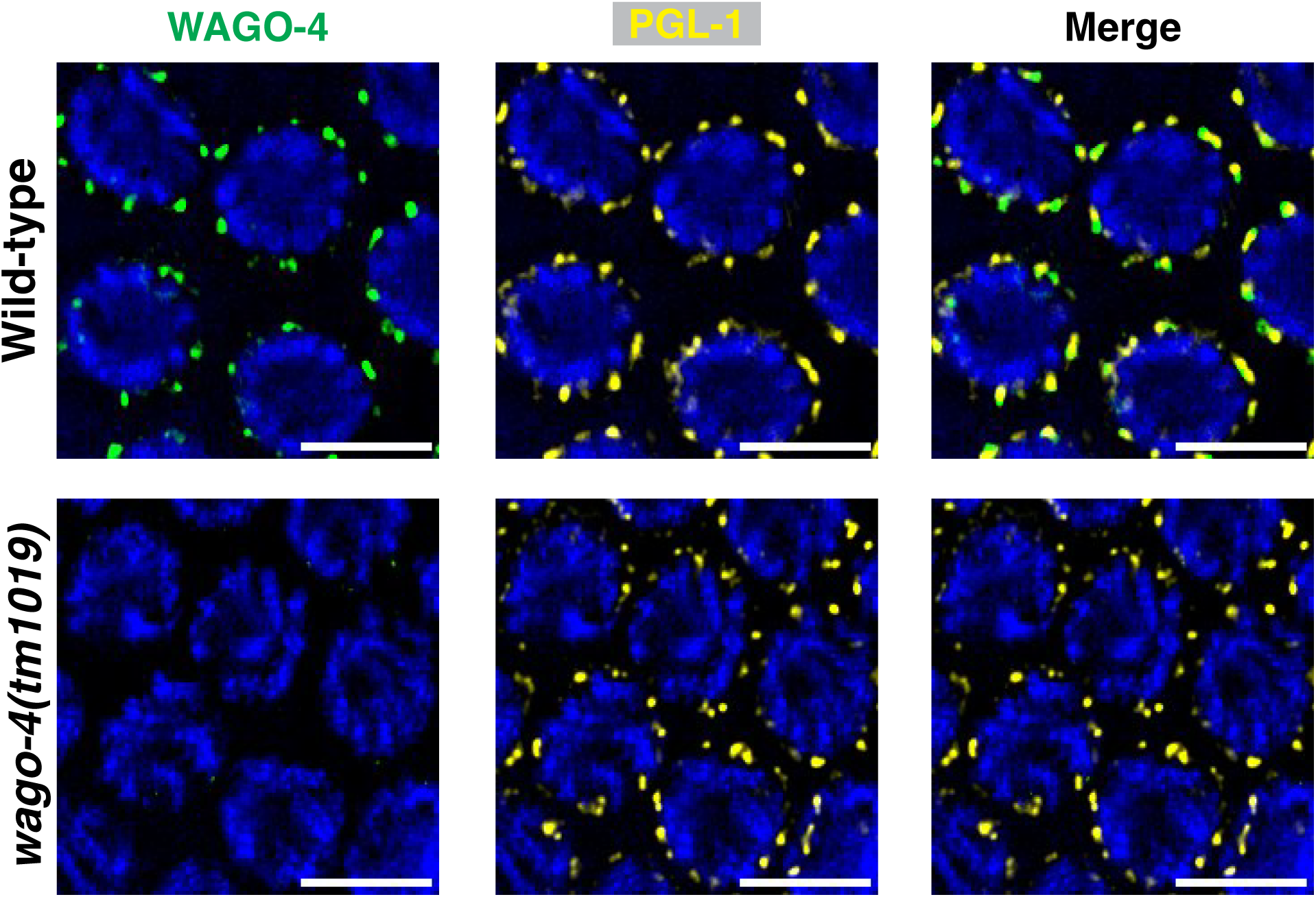
Immunofluorescent confirmation of WAGO-4 anti-guinea pig antibody specificity. Representative images of oocyte nuclei from wild-type (top) and *wago-4(tm1019)* during LP stained for WAGO-4 (green) and PGL-1 (yellow). Scale bars represent 1μm.

**Supplemental Figure 7:**
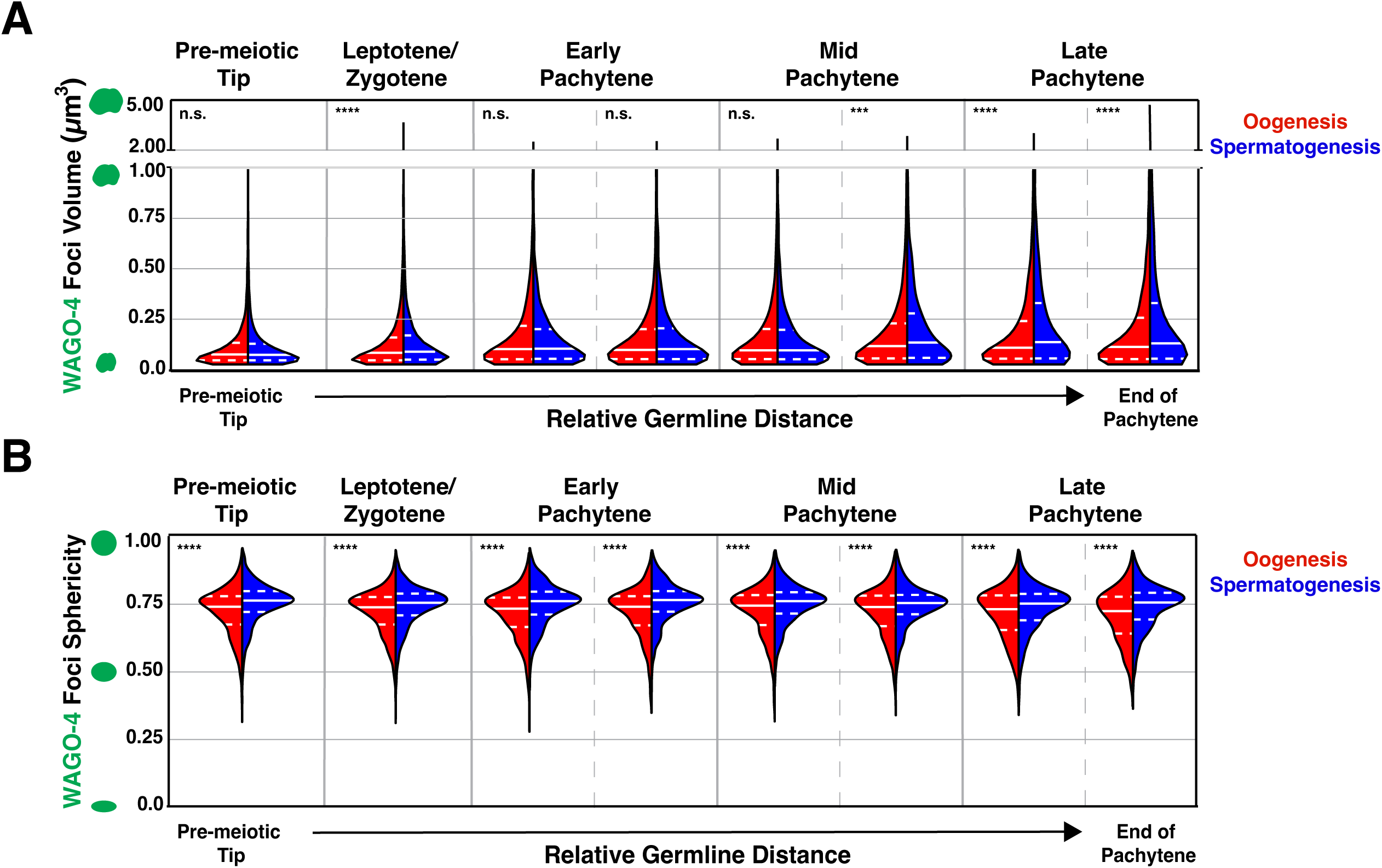
Volume and sphericity distribution of WAGO-4 foci during germ cell development. **(A)** Split violin plot depicting the volume distribution of WAGO-4 foci during oogenesis (red) and spermatogenesis (blue). **(B)** Split violin plot depicting the sphericity distribution of WAGO-4 foci during oogenesis (red) and spermatogenesis (blue). Dashed white lines represent the first and third quartiles, solid white line represents the second quartile/median. For all statistical tests, p-values were calculated using two-sample Kolmogorov-Smirnov test. n.s. p>0.05, *p<0.05, **p<0.01, ***p<0.001, ****p<0.0001.

## Notes

### Competing Interest Statement

The authors have declared no competing interest.

